# Termination of pre-mRNA splicing requires that the ATPase and RNA unwindase Prp43 acts on the catalytic snRNA U6

**DOI:** 10.1101/659029

**Authors:** Rebecca Toroney, Klaus H. Nielsen, Jonathan P. Staley

**Affiliations:** Moleculera Labs, Oklahoma City, OK, 73104, USA; Department of Molecular Genetics and Cell Biology, University of Chicago, IL 60637, USA

**Keywords:** pre-mRNA splicing, spliceosome, intron, disassembly, fidelity, U6 snRNA, Prp43, DEAH box, ATPase, helicase

## Abstract

The termination of pre-mRNA splicing functions to discard suboptimal substrates, thereby enhancing fidelity, and to release excised introns in a manner coupled to spliceosome disassembly, thereby allowing recycling. The mechanism of termination, including the RNA target of the DEAH-box ATPase Prp43, remains ambiguous. We discovered a critical role for nucleotides at the 3’-end of the catalytic U6 small nuclear RNA in splicing termination. Though conserved sequence at the 3’-end is not required, 2’ hydroxyls are, paralleling requirements for Prp43 biochemical activities. While the 3’-end of U6 is not required for recruiting Prp43 to the spliceosome, the 3’ end crosslinks directly to Prp43 in an RNA-dependent manner. Our data indicate a mechanism of splicing termination in which Prp43 translocates along U6 from the 3’ end to disassemble the spliceosome and thereby release suboptimal substrates or excised introns. This mechanism reveals that the spliceosome becomes primed for termination at the same stage it becomes activated for catalysis, implying a requirement for stringent control of spliceosome activity within the cell.

## Introduction

Eukaryotic gene expression requires pre-mRNA splicing to excise introns, a process that is frequently targeted for regulation and often perturbed in disease (Scotti and Swanson 2016; Fu and Ares 2014). An intron is excised from pre-mRNA in two steps: first, an intronic, branch site adenosine attacks the 5’ splice site, yielding a free 5’ exon and a characteristic branched lariat intermediate; second, the free 5’ exon attacks the 3’ splice site, excising the lariat intron and ligating the exons. Conserved intronic sequences define these three reactive sites of the substrate and recruit the catalyst of splicing – the spliceosome, which is conserved from budding yeast to humans and composed of 5 small nuclear RNAs (snRNAs) and over 80 proteins, as recently defined structurally by cryoelectron microscopy (Will and Lührmann 2011; Fica and Nagai 2017; Yan et al. 2019). Assembly of the spliceosome is coupled to intron recognition, resulting in a highly dynamic ribonucleoprotein machine (Staley and Guthrie 1998). Once splicing is complete, this coupling necessitates that splicing termination proceeds through disassembly of the spliceosome to release the excised intron product and also to recycle spliceosome components. Additionally, disassembly plays a central role in the fidelity of splicing (see below). Nevertheless, the mechanism of spliceosome disassembly remains under investigation.

Spliceosome assembly begins when the U1 small nuclear ribonucleoprotein (snRNP) complex binds the 5’ splice site and follows with the binding of the U2 snRNP to the branch site and then the recruitment of the U4/U6.U5 tri-snRNP (Will and Lührmann 2011). U1 snRNA is then replaced by U6 snRNA at the 5’ splice site, and U6 is liberated from a base pairing interaction with U4 snRNA (Staley and Guthrie 1999; Raghunathan and Guthrie 1998). These rearrangements enable U6 to form catalytic structures including a helix between U2 snRNA and U6 that juxtaposes the 5’ splice site and branch site for the branching reaction (U2/U6 helix I) and the intramolecular stem loop of U6 (the U6 ISL), which positions catalytic metals (Fica et al. 2013; Hilliker and Staley 2004; Madhani and Guthrie 1992; Sun and Manley 1995; Hang et al. 2015); U2 and U6 are also held together by a nearby helix – U2/U6 helix II (Fig. 1A) (Madhani and Guthrie 1994; Hang et al. 2015). Following release of U4 and a suite of proteins, the nineteen complex (NTC) and other factors bind, and then catalytic core of the spliceosome engages the substrate and catalyzes the splicing reactions (Galej et al. 2016; Fica et al. 2017; Liu et al. 2017; Fabrizio et al. 2009; Bai et al. 2017; Wilkinson et al. 2017; Wan et al. 2019; Yan et al. 2017; Wan et al. 2016). Finally, the mRNA is released from the spliceosome, and then the excised intron is released for degradation in a manner coupled to spliceosome disassembly (Company et al. 1991; Wagner et al. 1998; Schwer and Gross 1998; Arenas and Abelson 1997; Martin et al. 2002).

**Figure 1.**
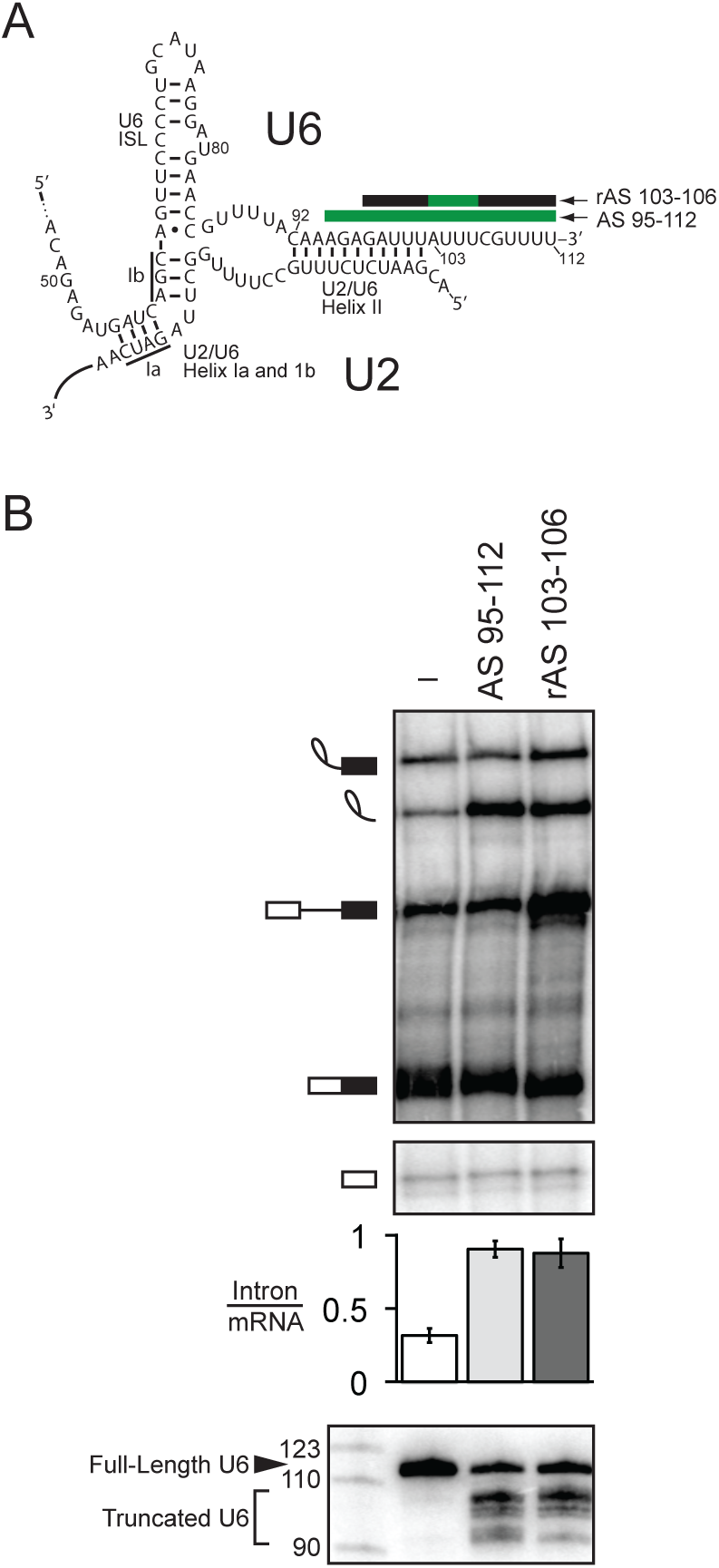
The very 3’ end of U6 is required for turnover of excised intron. **A**. Secondary structure of U6 and U2/U6 basepairing interactions at the catalytic core of the spliceosome. Anti-sense oligos used for RNase H cleavage are depicted by bars above their target U6 sequences; green indicates DNA nucleotides in the anti-sense oligo, while black indicates 2’-O-methyl nucleotides. **B**. Truncation of the 3’ end of U6 by ∼10 nts precludes turnover of the excised intron. Denaturing PAGE analysis of radiolabeled *ACT1* pre-mRNA following *in vitro* splicing in yeast extracts (yJPS860) that were first subjected to RNase H cleavage to truncate the 3’ end of U6 snRNA, directed by oligonucleotides depicted in **A**. Quantitation of intron turnover for each reaction was calculated as the molar ratio of excised intron to mRNA and is shown below the gel and is represented as the mean ± SD for three independent replicates. Cleavage of U6 was monitored by northern blot (bottom panel) with a radioactive probe directed to nucleotides 28-54 of U6.

Many of the RNA rearrangements and protein dynamics characteristic of the spliceosome are driven by members of the SF2 superfamily of nucleic acid-dependent ATPases, including eight members that are conserved from budding yeast to humans (Cordin and Beggs 2013). These ATPases share a common core of two RecA-like domains with over ten conserved motifs that function in RNA binding, ATP binding and hydrolysis, and in coupling of RNA binding to the ATPase cycle (Fairman-Williams et al. 2010; Ozgur et al. 2015). These motifs define four families that act on RNA – the DEAD family and three families that are collectively called the DExH group of families and include the Ski2-like family, the DEAH/RHA family, and the viral NS3/NPH-II family. Whereas the DEAD family binds single stranded RNA (ssRNA) in an ATP dependent manner and unwinds RNA by binding a strand of a duplex and distorting the duplex, the DExH families bind ssRNA in an ATP-independent manner and are thought to unwind by translocating 3’ to 5’ along ssRNA in an ATP-dependent manner. In this way, DExH families either plow directly through an RNA duplex or RNP complex (Pyle 2008) or pull on ssRNA to disrupt an RNA duplex or RNP complex at a distance (Semlow et al. 2016). Of the eight SF2 ATPases that play a conserved role in splicing, the three DEAD box ATPases function early in assembly; the Ski2-like DExH ATPase functions during spliceosome activation; and the four DEAH ATPases function at the catalytic stage of splicing, suggesting distinct activities are required at different stages of splicing (Cordin and Beggs 2013). All four DEAH ATPases impact the interaction of the pre-mRNA substrate with the spliceosome: after spliceosome activation, Prp2 enables positioning of the branch site for branching (Wlodaver and Staley 2014; Warkocki et al. 2015; Krishnan et al. 2013; Liu and Cheng 2012; Rauhut et al. 2016; Yan et al. 2016); after branching, Prp16 promotes repositioning of the branch site to enable 3’ splice site docking for exon ligation (Ohrt et al. 2013; Galej et al. 2016; Wan et al. 2016; Semlow et al. 2016; Schwer and Guthrie 1992); after exon ligation, Prp22p triggers release of the mRNA (Company et al. 1991; Wilkinson et al. 2017; Bai et al. 2017; Liu et al. 2017; Wagner et al. 1998; Schwer and Gross 1998); and subsequently Prp43, which also functions in ribosome biogenesis (Leeds et al. 2006; Combs et al. 2006; Lebaron et al. 2005), releases the excised intron for decay and coordinately disassembles the spliceosome for recycling (Arenas and Abelson 1997; Martin et al. 2002; Wan et al. 2017; Zhang et al. 2019).

Importantly, a number of these spliceosomal SF2 ATPases not only promote splicing of optimal substrates but also antagonize splicing of suboptimal substrates, thereby enhancing the specificity of splicing (Semlow and Staley 2012). The spliceosome utilizes these ATPases, after the spliceosome binds a substrate, to enable a second inspection of a substrate. For example, the specificity of the branching and exon ligation reactions is enhanced by the DEAH ATPases Prp16p and Prp22p, respectively; whereas both of these ATPases promote optimal substrates after branching and exon ligation, respectively, they also function to antagonize suboptimal substrates by competing with these chemical reactions in a manner thought to establish kinetic proofreading (Mayas et al. 2006; Koodathingal et al. 2010; Burgess and Guthrie 1993). Importantly, these rejection activities in splicing are effectively reversible in that they allow the spliceosome, after rejecting one potential splicing site, to select an alternative site (Semlow et al. 2016). However, in cases where the spliceosome fails to ever engage an optimal site, the spliceosome discards a substrate via the action of the DEAH ATPase Prp43p (Pandit et al. 2006; Mayas et al. 2010; Koodathingal et al. 2010; Chen et al. 2013). Because Prp43p functions to discard substrates at multiple stages of splicing, in addition to disassembling the spliceosome at the end of a splicing cycle (Arenas and Abelson 1997; Martin et al. 2002), Prp43p functions as a general terminator of splicing. Through its discard activity, Prp43p functions to repress cryptic 3’ splice sites and thereby promote fidelity (Mayas et al. 2010). Interestingly, the roles of Prp22p and Prp43p in rejecting and discarding suboptimal substrates have been repurposed in a range of ascomycetes fungi for the biogenesis of telomerase RNA, which corresponds to a released 5’ exon intermediate in these species (Kannan et al. 2013; 2015; Qi et al. 2015).

As for the other spliceosomal DEAH ATPases (Kim et al. 1992; Wagner et al. 1998; Schwer and Guthrie 1991), the ATPase activity of Prp43p is stimulated by RNA but not DNA, and Prp43p only binds RNA (Tanaka and Schwer 2006; Martin et al. 2002). Consistent with its membership in the DExH family of ATPases, Prp43p unwinds duplexes specifically with a 3’ overhang (He et al. 2017), implying translocation along RNA in the 3’ to 5’ direction (Jankowsky 2011). Crystal structures of Prp43, Prp22, as well as of the DEAH ATPase MLE, bound to RNA and non-hydrolyzable ATP provided the first structural insights into the mechanism of translocation by the DEAH family members (He et al. 2010; Prabu et al. 2015; Tauchert et al. 2017; He et al. 2017; Hamann et al. 2019; Chen et al. 2018). The two RecA domains bind an RNA stack of four nucleotides that is bookended at the 5’ end by an elongated beta-hairpin, as in the viral SF2 ATPase NS3 (Luo et al. 2008; Gu and Rice 2010; Appleby et al. 2011), and at the 3’ end by a DEAH-specific elongation of motif Ib, which includes a residue that is required for RNA unwinding and mutated in human Prp43p in acute myeloid leukemias (Faber et al. 2016). A comparison of Prp43p structures in distinct nucleotide states has revealed a specific conformational change – rearrangement of motif Va in the RecA2 domain, as observed in NS3, a rearrangement anticipated to impact RNA binding, suggesting that this motif functions to couple the ATP cycle to RNA binding and translocation (He et al. 2017; Appleby et al. 2011; Gu and Rice 2010; Hamann et al. 2019). Altogether, structures of the DExH proteins suggest a fundamental mechanism for translocation in which the RecA domains inch along RNA, one nucleotide at a time.

Prp43p requires a co-factor of the G-patch protein family (Aravind and Koonin 1999; Robert-Paganin et al. 2015) for efficient ATPase and RNA unwinding activity. Indeed, distinct G-patch proteins serve to activate Prp43p in different processes, such as splicing and ribosome biogenesis (Tanaka et al. 2007; Lebaron et al. 2009; Chen et al. 2014b; Heininger et al. 2016; Boon et al. 2006; Pandit et al. 2006; Tsai et al. 2005). In splicing, the conserved G-patch protein Ntr1p/Spp382p activates Prp43p and forms the NTR (<u>NT</u>C-<u>R</u>elated) complex with Prp43p, Cwc23p, and Ntr2p, a factor that is found primarily in fungi and plants. Ntr1p, Cwc23p, and Ntr2p help to recruit Prp43p to the spliceosome (Tsai et al. 2005). The regulation of this recruitment is important, given that Prp43p acts as a general terminator of splicing (Koodathingal et al. 2010; Mayas et al. 2010; Pandit et al. 2006; Arenas and Abelson 1997; Martin et al. 2002; Chen et al. 2013). Ntr2p and/or a portion of Ntr1p appear to enforce this regulation, because the G-patch domain of Ntr1p alone in conjunction with Prp43p enables disassembly of spliceosomal complexes otherwise refractory to termination (Fourmann et al. 2016; 2017).

As for many members of the SF2 family of ATPases, a long-term goal has been to determine the physiological, RNA target of Prp43p. Crosslinking *in vivo* has revealed that Prp43p interacts directly with rRNA and snoRNA as well as U6 snRNA, implicating these RNAs as targets in ribosome biogenesis and splicing, respectively (Bohnsack et al. 2009). However, a recent *in vitro* study has implicated the U2-branch site duplex as the target for Prp43p in splicing (Fourmann et al. 2016). A structure of the budding yeast spliceosome poised for disassembly is consistent with Prp43p targeting either U6 or the branch site duplex (Wan et al. 2017), whereas a structure of the analogous human spliceosomal intermediate favors U6 as the target (Zhang et al. 2019).

In this study, we have gained insight into the mechanism of Prp43p action through an investigation of the function of the 3’ end of U6. We have found that the 3’ end of U6 is required for the final stage of splicing – spliceosome disassembly and intron release – and also for the fidelity of exon ligation, through a requirement in the discard of a rejected, suboptimal substrate. These functions of the 3’ end of U6 parallel the functions of Prp43 (Arenas and Abelson 1997; Mayas et al. 2010; Martin et al. 2002; Chen et al. 2013). Further, the requirements for the 3’ end of U6 in intron turnover parallel the requirements for Prp43p – both require RNA, because DNA is insufficient, and neither requires a specific sequence, including the conserved sequence of the 3’ end of U6 (Tanaka and Schwer 2006; Martin et al. 2002). Indeed, Prp43p crosslinks *in vitro* to the 3’ end of U6. Our observations imply a mechanism for spliceosome disassembly and intron release in which Prp43p acts as a winch that pulls on the 3’ end of U6 to disrupt interactions between the catalytic U6 snRNA and the spliceosome.

## Results

### The 3’ end of U6 is required for excised intron release and spliceosome disassembly

*In vivo*, the 3’ end of U6 is essential for growth in yeast and plays a critical role in stabilizing U6 snRNA through the recruitment of the La homolog protein Lhp1p and subsequently the Lsm2-8 complex. Unlike Lhp1p, the Lsm complex tolerates the terminal, cyclic 2’, 3’ phosphate of U6 in metazoans and the terminal 3’ monophosphate in yeast, both of which result from 3’-end maturation during U6 biogenesis; whereas Lhp1 binds the UUU_OH_ terminus of all RNA polymerase III transcripts (Wolin and Cedervall 2002), the Lsm complex binds primarily to the GUUUU terminal sequence of U6 (Pannone et al. 2001; Achsel et al. 1999; Bordonne and Guthrie 1992; Wolff and Bindereif 1995; Zhou et al. 2014; Licht et al. 2008; Montemayor et al. 2018). The 3’ end of U6 has additionally been shown *in vitro* to play an early role in splicing by promoting the annealing of U6 to U4, again through recruitment of the Lsm complex and also through the subsequent recruitment of Prp24p (Ryan et al. 2002; Vidal et al. 1999; Rader and Guthrie 2002; Licht et al. 2008). Surprisingly, the 3’ end of U6 has also been implicated *in vitro* in playing a late role in splicing – in the turnover of the excised lariat intron, long after Prp24p has dissociated from U6 during U4/U6 annealing, and long after the Lsm2-8 complex has dissociated during spliceosome activation. Specifically, in whole cell extract from budding yeast, either reconstituting extract with 3’ end-truncated U6 or targeting the 3’ end of U6 for RNase H-mediated cleavage with a DNA oligonucleotide complementary to the last 31 nts of U6 both stabilized the excised, lariat intron product of splicing, preventing its turnover in such reactions (Ryan et al. 2002; Fabrizio et al. 1989). Truncation of the 3’ end of U6 by as few as 23 nts (nts 90-112) was sufficient to stabilize the excised intron (Ryan et al. 2002).This region of U6 participates in two structures – U2/U6 helix II (nts 92-102) and the essential component of the intramolecular telestem (nts 92-95) – and also binds to the Lsm2-8 complex (nts 108-112).

To distinguish more finely the region of U6 essential for intron turnover, we targeted RNase H cleavage even closer to the 3’ end of U6 in budding yeast extract, using an oligonucleotide complementary to the last 18 nts of U6 (AS 95-112) or a chimeric DNA/2’-O-methyl oligonucleotide (rAS 103-106) that directed RNase H to cleave 7-10 nts from the 3’ end (Fig. 1A). Both oligonucleotides resulted in a substantial accumulation of excised, lariat intron, as reflected by an increase in the molar ratio of excised intron to mRNA from 0.32 ±0.05 to 0.91 ±0.06 or to 0.88 ±0.10, respectively (Fig. 1B). In both cases, the native U6 was truncated predominantly to nt 103 or 104 at the 3’ end of U2/U6 helix II, presumably through the action of a 3’ exonuclease (cf. (Ryan et al. 2002)). Thus, we conclude that the last ∼10 nts of U6 are essential for turnover of the excised, lariat intron, a region outside of U2/U6 helix II and the U6 telestem but including the binding site for the Lsm2-8 complex.

Turnover of the excised, lariat intron by exonucleases requires debranching, which requires release from the spliceosome (Martin et al. 2002). The spliceosome releases the excised, lariat intron in two ATP-dependent steps. First, the DEAH ATPase Prp22p releases the mRNA (Schwer and Gross 1998; Company et al. 1991; Wagner et al. 1998), which then allows the DEAH ATPase Prp43p to subsequently disassemble the spliceosome and release the excised lariat intron (Arenas and Abelson 1997; Martin et al. 2002). To determine the stage at which the 3’ end of U6 is required for excised intron turnover, we analyzed splicing reactions by glycerol gradients, which separate released splicing products from spliceosome-bound products. In splicing reactions in which U6 was cleaved with AS 90-112, the stabilized, excised lariat intron did not migrate at the top of the gradient, demonstrating that the 3’ end of U6 does not somehow impact debranching directly, after release of the intron from the spliceosome. Instead, the stabilized, excised lariat intron co-migrated with spliceosomes deep in the gradient, implicating a direct or upstream role for the 3’ end of U6 in release of the intron from the spliceosome (Supplemental Fig. S1A, B).

To account fully for the excised, lariat intron released from the spliceosome, we precluded turnover of these lariats by assembling splicing reactions in *dbr1Δ* extract lacking the debranchase Dbr1p. In these extracts, cleavage of the 3’ end of U6 shifted the migration of the excised lariat intron from shallow fractions of the gradient to deep, spliceosomal fractions, establishing that the 3’ end of U6 is required for release of the excised lariat intron from the spliceosome (Fig. 2A, cf. first and second panels; Fig. 2B, right panel); northern blotting showed that truncated U6 also comigrates with the spliceosome deep in the gradient, confirming that U6 lacking ∼10 nts at the 3’ end incorporates into spliceosomes (Supplemental Fig. S1C). This requirement of the 3’ end of U6 in excised intron release parallels the requirement for Prp22p and Prp43p in intron release; specifically, intron release was similarly impeded by adding to splicing reactions either recombinant Prp22p having a dominant-negative K512A mutation or recombinant Prp43p having a dominant-negative Q423E mutation, as expected (Leeds et al. 2006; Schwer and Gross 1998) (Fig. 2A, compare the first panel with the third and fourth panels; Fig. 2B, right panel). Cleavage of the 3’ end of U6 did not shift the migration of mRNA, in contrast to the excised intron, from shallow fractions to spliceosome-containing fractions, indicating that the 3’ end of U6 is not required for mRNA release (Fig. 2A, B). This lack of a requirement for the 3’ end of U6 in mRNA release contrasts with a requirement for Prp22p in mRNA release and parallels the lack of a requirement for Prp43p; indeed, whereas the Prp22p mutation retained mRNA in spliceosome-containing fractions, the Prp43p mutation did not, as expected (Fig. 2A,B). These data therefore demonstrate that the 3’ end of U6 is required after mRNA release, specifically at the stage of intron release and spliceosome disassembly; these data thus implicate components of the NTR complex, including Prp43p, as factors that mediate the requirement for the 3’ end of U6 in spliceosome disassembly (Small et al. 2006).

**Figure 2.**
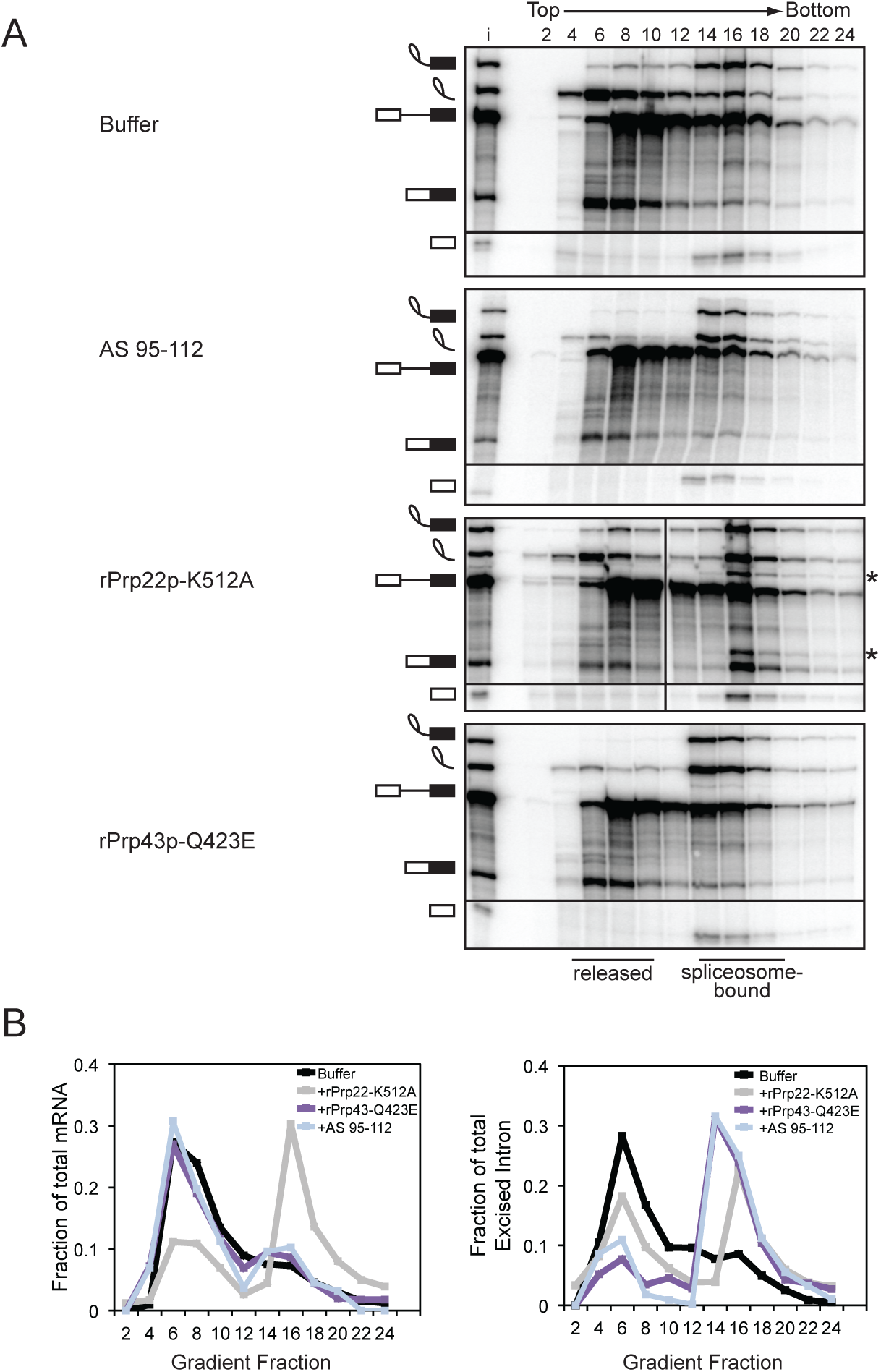
The 3’ end of U6 is required for release of the excised intron. **A**. Truncation of the 3’ end of U6 by ∼10 nts precludes release of the excised intron from the spliceosome. Radiolabeled *ACT1* pre-mRNA was spliced in mutant *dbr1Δ* (yJPS799) extracts subjected to RNase H cleavage directed by DNA oligo AS 95-112 (see Fig. 1A) or supplemented with either buffer, mutated rPrp22p-K512A, or mutated rPrp43p-Q423E. Splicing reactions were fractionated on a glycerol gradient; input (i) and fraction numbers are indicated above the top panel. Fractions containing released or spliceosome-bound splicing species are highlighted below the bottom panel. The dividing line in the rPrp22p-K512A panel indicates an empty lane that was removed for consistency between panels. The asterisks identify excised intron and mRNA resulting from splicing at an upstream, suboptimal 3’ splice site due to a loss of fidelity resulting from the Prp22p-K512A mutant (Mayas et al. 2006). See also Figure S1. **B**. Quantitation of mRNA (left) or excised intron (right) from gradients in **A**. Data are normalized as a fraction of total mRNA or total excised intron within each gradient.

### The 3’ end of U6 is required for fidelity and discard of a suboptimal lariat intermediate

In addition to its role during canonical disassembly following exon ligation, Prp43p – likely in the context of the NTR complex (Chen et al. 2013; Su et al. 2018) – also contributes to splicing fidelity by promoting, through spliceosome disassembly, discard of suboptimal substrates such as pre-mRNA rejected by Prp16p and intermediates rejected by Prp22p (Koodathingal et al. 2010; Mayas et al. 2010). Thus, if the requirement for the 3’ end of U6 in spliceosome disassembly and release of a canonical intron reflects a requirement for Prp43p, then the 3’ end of U6 should also be required for Prp43p-dependent discard of suboptimal substrates and splicing fidelity. To test these predictions, we first assayed for the requirement of the 3’ end of U6 in the discard of splicing intermediates after Prp22p-mediated rejection of a suboptimal 3’ splice site, again by glycerol gradient analysis of *in vitro* splicing reactions in *dbr1Δ* extract. Specifically, we utilized a *UBC4* pre-mRNA containing a suboptimal, UgG 3’ splice site, which is rejected by Prp22p before the stage of exon ligation and then discarded as 5’ exon and lariat intermediate by Prp43p (Mayas et al. 2010; 2006). As expected, with full length U6 a substantial fraction of the rejected and discarded UgG lariat intermediate migrated in fractions near the top of the gradient (Fig. 3A, top panel; Fig. 3B). By contrast, with truncated U6, the UgG lariat intermediate migrated primarily in deeper, spliceosome-containing fractions, indicating a defect in discard of the rejected intermediate (Fig. 3A, bottom panel; Fig. 3B). This defect parallels an equivalent defect conferred by the Q423E mutation in Prp43p (Fig. 3A, middle panel; Fig. 3B) (Mayas et al. 2010). Thus, like Prp43, the 3’ end of U6 is required for discard of a suboptimal intermediate.

**Figure 3.**
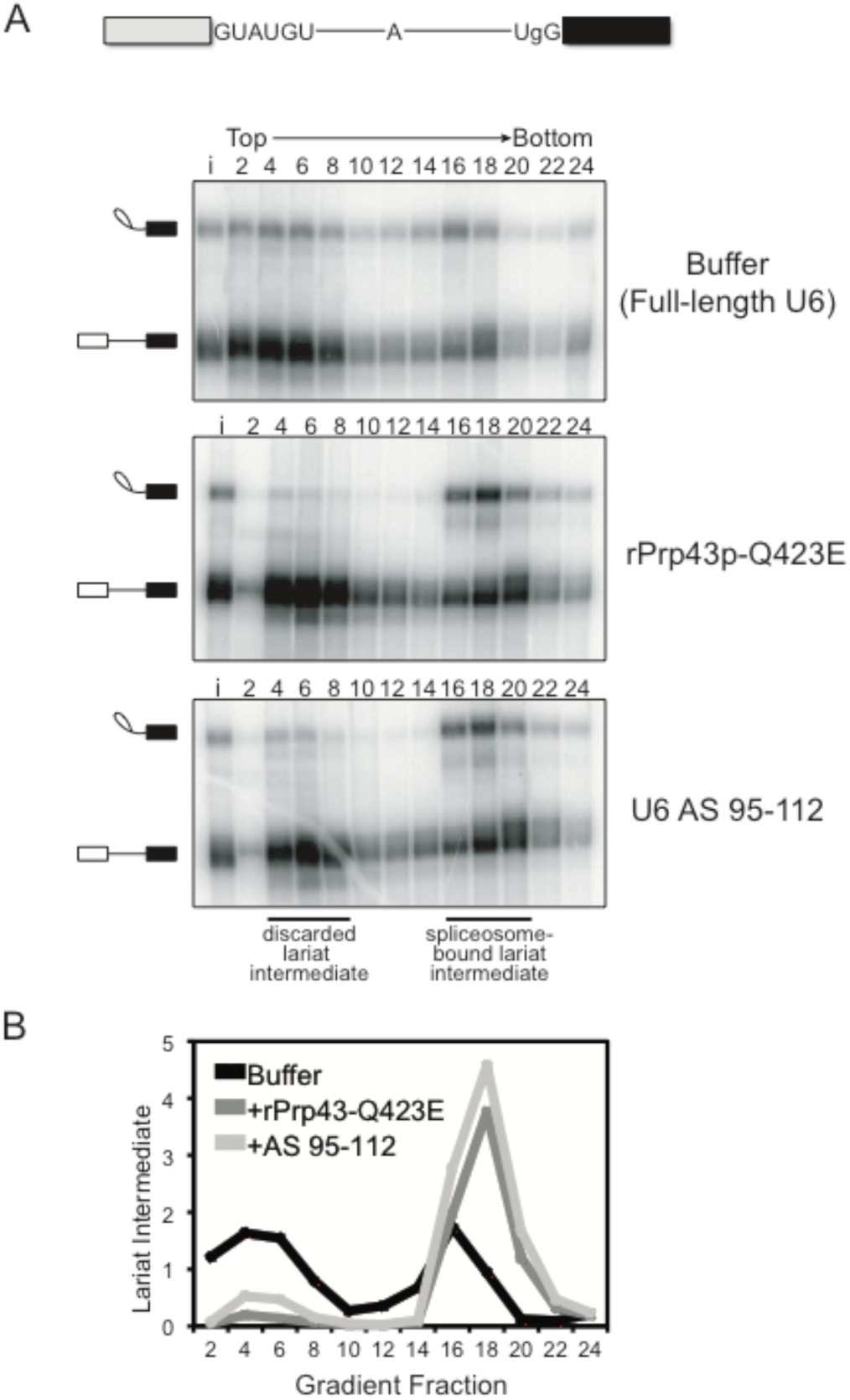
The 3’ end of U6 is required *in vitro* for Prp43p-dependent discard of a suboptimal substrate. **A**. As with the rPrp43p-Q423E mutant (Mayas et al. 2010), 3’-truncated U6 impedes discard of a suboptimal lariat intermediate. Radiolabeled *UBC4* pre-mRNA having a suboptimal UgG 3’ splice site was spliced in mutant *dbr1Δ* (yJPS799) extracts with buffer, with rPrp43p-Q423E or after RNase H cleavage of U6 in extract with DNA oligo AS 95-112 (see Figure 1A). Splicing reactions were fractionated on a glycerol gradient; input (i) and fraction numbers are indicated above the top panel. The migration of spliceosome-bound and discarded lariat intermediate is indicated. See also Figure S2. **B**. Quantitation of lariat intermediate levels from glycerol gradients in **A**. Data are normalized to input levels of lariat intermediate.

Through its role in discard, Prp43p promotes the fidelity of splicing by minimizing formation of cryptic mRNAs at the stage of exon ligation (Mayas et al. 2010). For example, with a *UBC4* pre-mRNA having a UgG 3’ splice site mutation, Prp43p not only discards the suboptimal lariat intermediate but also minimizes splicing at a cryptic 3’ splice site, 6 nucleotides upstream of the suboptimal, UgG 3’ splice site (Supplemental Fig. S2). We found that in *DBR1* wild-type extract, this proofreading function of Prp43p was also dependent on the 3’ end of U6. Specifically, truncation of U6 with oligonucleotide AS 95-112 led to stabilization of the suboptimal lariat intermediate and increased production of cryptic mRNA from the UgG substrate, comparable to the increase conferred by recombinant mutant Prp43p-Q423E (Supplemental Fig. S2). Thus, a requirement for the 3’ end of U6 again parallels a requirement for Prp43p. In sum, these data illustrate a striking correlation between the requirement for Prp43p and the 3’ end of U6 in disassembly, discard, and fidelity.

### The 3’ end of U6 is required after binding of the NTR complex

Given the strong correlation between the requirements for Prp43p and the 3’ end of U6, we investigated the mechanism by which the 3’ end of U6 functions in spliceosome disassembly and intron release by first determining whether nucleotides at the 3’ end of U6 are required before or after recruitment of the NTR complex to the spliceosome, following release of mRNA by Prp22p. Specifically, we stalled splicing of *ACT1* pre-mRNA at the intron release stage by cleavage of U6 with oligonucleotide AS 95-112 or with dominant-negative rPrp43p-Q423E, as a positive control for NTR complex association (Small et al. 2006), or with dominant-negative rPrp22p-K512A, as a negative control for NTR complex association (Chen et al. 2013). We then tested for binding of the NTR complex to spliceosomes by assaying for co-immunoprecipitation of excised intron with the NTR complex, using antibodies to Ntr1p, Ntr2p, or Prp43p (Tanaka et al. 2007; James et al. 2002). When U6 was truncated, immunoprecipitation of Ntr1p, Ntr2p, or Prp43p enriched for excised intron relative to other splicing species, especially pre-mRNA and mRNA (Fig. 4, top panel, cf. lane 3 to lanes 6, 9, and 11), as observed for the immunoprecipitation of Ntr1p and Ntr2p in reactions stalled by rPrp43p-Q423E (Fig. 4, cf. lanes 2, 5, and 8); by contrast, such enrichment was not observed in reactions stalled by rPrp22p-K512A, despite the accumulation of comparable levels of excised intron (Fig. 4, cf. lanes 4, 7, 10, and 12, respectively). For splicing reactions stalled by the truncation of U6, we confirmed by northern that immunoprecipitation of the NTR complex not only co-immunoprecipitated excised intron but also truncated U6 (Fig. 4, bottom panel, lanes 6, 9, 11). These data indicate that the 3’ end of U6 is not required for the recruitment of the NTR complex, indicating this region of U6 plays a role downstream of NTR binding to promote intron release and spliceosome disassembly; note that whereas Cwc23p has also been shown to complex with Ntr1p and Ntr2p, Cwc23p is not required for binding of Ntr1p and Ntr2p to intron-lariat spliceosomes or for release of excised intron (Su et al. 2018; Pandit et al. 2006), so the 3’ end of U6 cannot play a role in recruiting Cwc23p to promote spliceosome disassembly.

**Figure 4.**
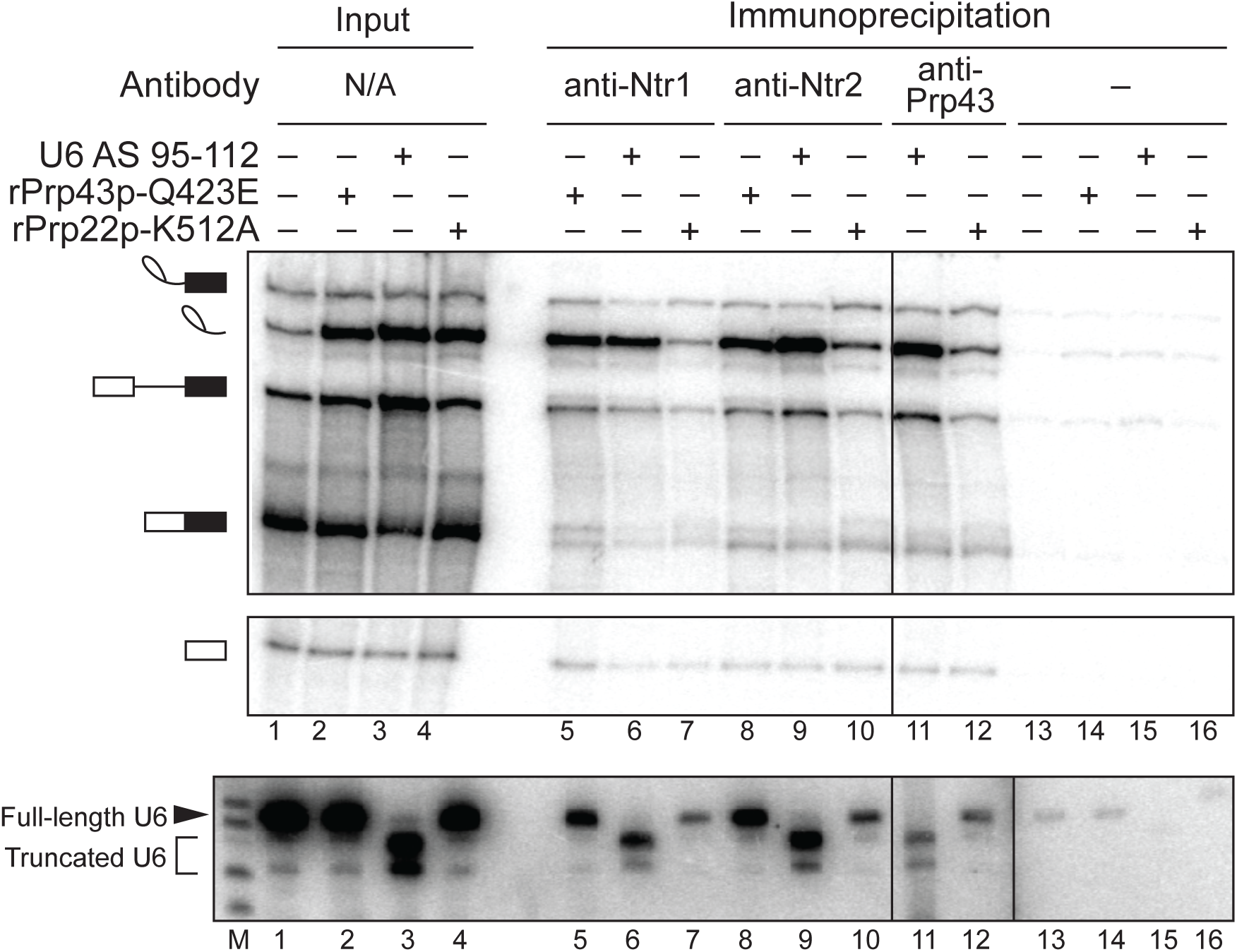
The 3’ end of U6 is required after the binding of Prp43p, Ntr1p, and Ntr2p to the spliceosome. Denaturing PAGE analysis of *in vitro* splicing reactions after immunoprecipitation with anti-Ntr1p, -Ntr2p, or -Prp43p antibodies (top, middle panels). Radiolabeled *ACT1* pre-mRNA was spliced in yeast extracts (yJPS1448) that were subjected to RNase H cleavage with DNA oligo AS 95-112. Immunoprecipitation was not performed with anti-Prp43p antibodies on reactions supplemented with rPrp43p-Q423E due to complicating effects of excess recombinant protein. Twenty percent of each reaction was analyzed as input. Cleavage and coimmunoprecipitation of U6 was monitored by northern blot with a radioactive probe directed to nucleotides 28-54 of U6 (bottom panel); “M” indicates a marker lane. The vertical dividing line in all three panels indicates a control lane that was omitted for clarity; the additional vertical dividing line in northern blot indicates an extra marker lane that was omitted for clarity. See also Figure S3 for tests for association of other factors, Lsm3p and Prp24p, with the spliceosome during disassembly.

While the requirement for the 3’ end of U6 after NTR binding suggests the 3’ end may impact the function of Prp43p directly, at other stages of splicing the 3’ end interacts with the Lsm2-8 complex and Prp24p, so these factors could in principle play roles in spliceosome disassembly and intron release. Although Prp24p dissociates after U4/U6 annealing and the Lsm proteins dissociate after spliceosome activation (Jandrositz and Guthrie 1995; Chan et al. 2003), it is not entirely clear when these factors re-associate with U6 in the splicing cycle, and a recent crystal structure of a U6 snRNP complex suggested the possibility that Prp24p and by implication the Lsm2-8 complex may promote displacement of U2 from U6 during spliceosome disassembly (Montemayor et al. 2014). Thus, we tested for association of these factors with spliceosomes at the stage of disassembly and intron release by stalling spliceosomes with the rPrp43p-Q423E mutant and assaying for association of the Lsm complex or Prp24p by immunoprecipitation directly from splicing reactions (Supplemental Fig. S3). As expected, Lsm3p co-immunoprecipitated U4, U5, and U6, reflecting association with the tri-snRNP, under all conditions tested, and Lsm3p co-immunoprecipitated pre-mRNA, when stalled by low ATP at the stage of U4 release; by contrast, immunoprecipitation of Lsm3p did not enrich for excised intron relative to pre-mRNA or mRNA, as compared to a control without antibody, whereas immunoprecipitation of Ntr2p did (Supplemental Fig. S3A). Similarly, Prp24p co-immunoprecipitated U6 snRNA, as expected, but did not co-immunoprecipitate accumulated excised intron (Supplemental Fig. S3B). In contrast, Ntr1p of the NTR complex did co-immunoprecipitate accumulated excised intron (Supplemental Figs. S3B). While we cannot rule out occlusion of epitopes, these data provide no evidence that either the Lsm2-8 complex or Prp24p associate with spliceosomes poised for disassembly; supporting these findings, purified spliceosomes poised for disassembly have not revealed the presence of Prp24p or Lsm proteins by mass spectrometry or the requirement of trans-acting factors for disassembly after NTR association (Chen et al. 2014a; Fourmann et al. 2013; Small et al. 2006; Tsai et al. 2007; Wan et al. 2017; Zhang et al. 2019). Together, these data provide no evidence for a role of Prp24p or the Lsm2-8 complex in spliceosome disassembly.

### The conserved sequence of the 3’ end of U6 is not required for disassembly

Given the parallels between the requirements of Prp43p and the 3’ end of U6 in spliceosome disassembly and given the association of the NTR complex with spliceosomes stalled by truncated U6, we proceeded to test a model in which the 3’ end of U6 serves as a substrate of Prp43p. Because SF1 and SF2 ATPases bind fundamentally to the backbone of nucleic acid and generally without sequence specificity (Fairman-Williams et al. 2010) (with some exceptions, e.g. (Prabu et al. 2015)), this model predicts that the conserved sequence at the 3’ end of U6, required for Lsm2-8 complex binding (Zhou et al. 2014; Achsel et al. 1999; Vidal et al. 1999), is not required for spliceosome disassembly. To test this prediction, we mutated the U-rich sequence at the 3’ end of U6 to AG- or AC-rich sequences, calculated by *m*Fold (Zuker 2003) to form minimal to no secondary structure (100_AG and 100_AC; Fig. 5A). Because the three uracils at nts. 100-102 form basepairs with U2 as part of U2/U6 helix II, we also included a variant that did not alter these nucleotides (103_AG; Fig. 5A). Significantly, none of these sequence variants compromised intron turnover (Fig. 5B), establishing that spliceosome disassembly does not require a specific sequence at the 3’ end of U6, thus establishing functional evidence that intron turnover does not require the sequence-dependent Lsm2-8 complex (Achsel et al. 1999; Zhou et al. 2014). Moreover, these data are consistent with a role for Prp43p in acting on the 3’ end of U6.

**Figure 5.**
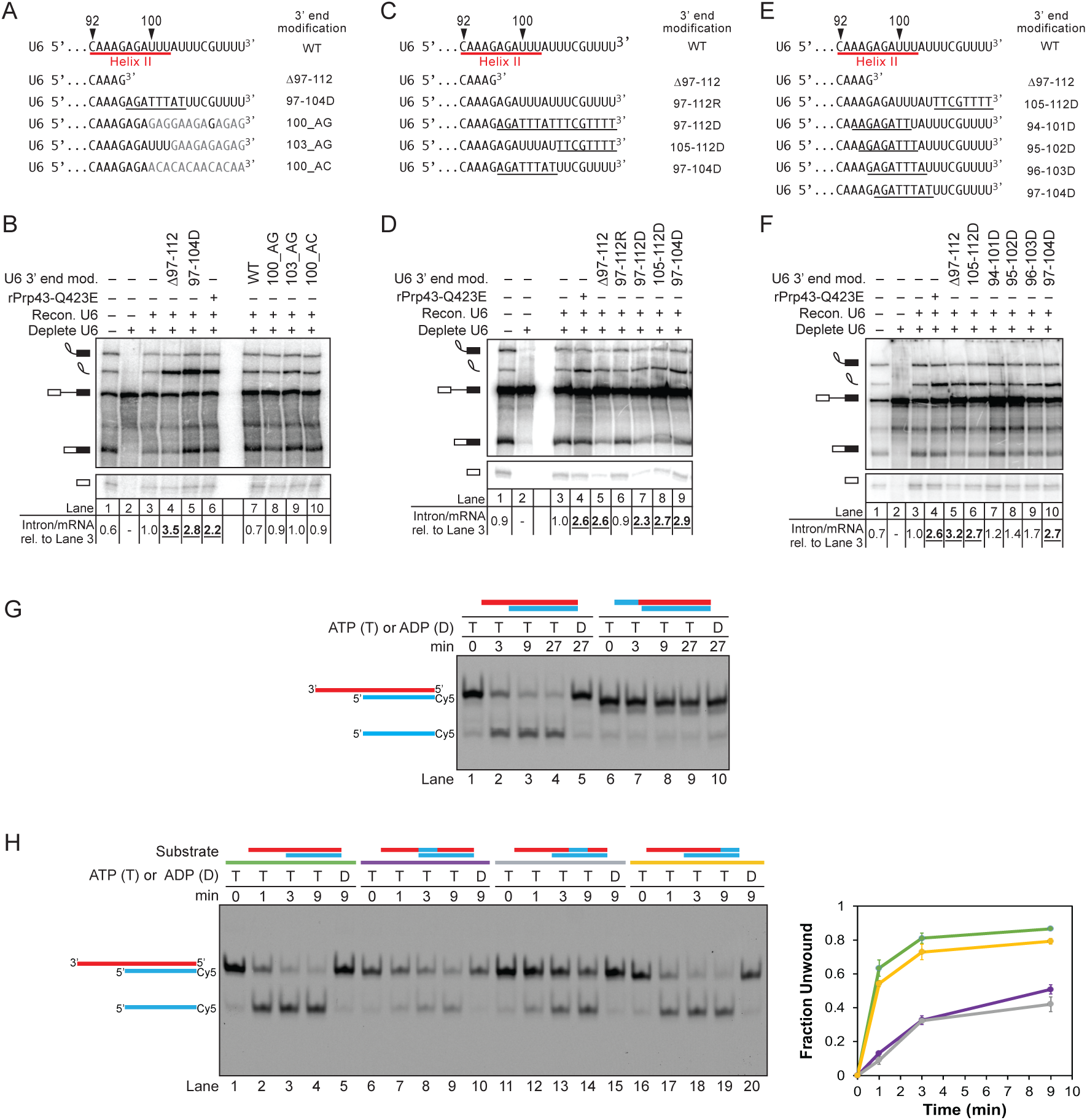
Spliceosome disassembly does not require the sequence of the 3’ end of U6 but does require the 2’ hydroxyls of the 3’ end. **A**. U6 3’ end sequences are depicted showing mutations used in B. “WT” corresponds to full-length, unmodified, transcribed U6 snRNA. U6 Δ97-112 and 97-112D were constructed by transcribing nucleotides 1-96 of U6 and then either using as is (Δ97-112) or first ligating to a DNA oligo (97-112D). The remaining variants were constructed by ligating segments of synthetic RNA. **B**. Disassembly does not require specific sequence of the 3’ end of U6. Denaturing PAGE analysis of radiolabeled *in vitro* splicing reactions using *ACT1* pre-mRNA in extracts (ySCC1) reconstituted (recon.) with the indicated U6 variants. U6 Δ97-112 and 97-112D correspond to positive controls that block disassembly and stabilize excised intron, as shown in D. Quantitation of intron turnover is shown below the gel, wherein the ratio of excised intron to mRNA was calculated and normalized relative to the wild-type U6 control in lane 3; values with a substantial increase of 2-fold or greater over wild-type are indicated in bold, underlined. **C**. U6 3’ end sequences are depicted showing deoxy substitutions used in D. Underlined bases are DNA. “WT” corresponds to full-length, unmodified, transcribed U6 snRNA; for the remaining U6 variants, the first 96 nucleotides were transcribed and then used as is (Δ97-112) or first ligated to synthetic oligonucleotides corresponding to unmodified RNA (97-112R), DNA (97-112D), or RNA-DNA chimeras (97-104D, 105-112D). **D**. Disassembly requires the 2’ hydroxyls of the 3’ end of U6. Denaturing PAGE analysis of splicing reactions using *ACT1* pre-mRNA in ySCC1 extracts reconstituted with the indicated U6 variants. See also Figures S4 and S5. **E**. U6 3’ end sequences are depicted showing deoxy substitutions used in F. U6 variants are shown and were constructed as in C. **F**. Disassembly only requires 2’ hydroxyls in the last ∼16 nts of the 3’ end of U6. Denaturing PAGE analysis of splicing reactions using *ACT1* pre-mRNA in extracts (yJPS866) reconstituted with the indicated U6 variants. **G**. Prp43p unwinds a duplex with a short RNA, but not DNA, single-stranded overhang *in vitro*. Unwinding of a 24 bp RNA/DNA duplex with a 10 nt single-stranded RNA overhang (red) or an 8 nt DNA substitution (blue) was assayed in the presence of rPrp43p, its activator rNtr1p (1-120), and ATP (T) or ADP (D), as a negative control. **H.** Prp43p-mediated unwinding is impeded by DNA at the proximal end and middle of a duplex but not at the distal end. Unwinding of a 24 bp RNA/DNA duplex with or without an 8 nt DNA substitution and with a 16 nt single-stranded RNA overhang was assayed in the presence of rPrp43p, its activator rNtr1p (1-120), and ATP (T) or ADP (D), as a negative control. Red indicates RNA; blue, DNA. Quantification is shown for the average of two replicate experiments; error bars indicate the range of values; unwinding was calculated as the fraction of released oligo relative to the total oligo – single-stranded and duplexed. Color coding in the left panel serves as a key for the right pane. In panels B, D, and F, “mod.” = “modification”.

### The 2’ hydroxyls of the 3’ end of U6 are required for spliceosome disassembly

Whereas Prp43p does not depend on a specific sequence for binding to nucleic acid, it does depend on the 2’ hydroxyls of RNA both for binding to model substrates and for nucleic acid-dependent ATPase activity (Tanaka and Schwer 2006; Martin et al. 2002). We therefore tested whether spliceosome disassembly similarly depends on the 2’ hydroxyls of RNA. Relative to U6 snRNA controls, U6 substituted in the last sixteen nucleotides with DNA (97-112D) accumulated excised intron relative to mRNA, similar to 3’ truncated U6 (Δ97-112) or the rPrp43p-Q423E mutant (Fig. 5C and D, cf. lane 3-7). This requirement for RNA at the 3’ end of U6 is also consistent with a role for Prp43p action at the 3’ end of U6.

To define precisely the region of U6 that must be RNA to promote intron turnover, we varied the length and position of DNA substitutions and assayed for intron turnover (Fig. 5C-F). Consistent with our finding that truncation of as few as 9 or 10 nucleotides from the 3’ end results in a defect in intron turnover (Fig. 1B), as few as 8 nucleotides of DNA at the 3’ end blocked intron turnover as robustly as rPrp43p-Q423E (Fig. 5C and D, cf. lanes 4 and 8); 6 or 4 nucleotides of DNA at the 3’ end of U6 were insufficient to similarly compromise intron turnover (Supplemental Fig. S4). Interestingly, placing the 8-nt. DNA block 8 nucleotides upstream of the 3’ end and overlapping with U2/U6 helix II (Fig. 5C and 5D, lane 9) still strongly inhibited intron turnover, indicating that 8 nts of RNA at the 3’ end of U6 are not sufficient for spliceosome disassembly. By contrast, 8-nt. DNA blocks 16, 20, or 34 nts upstream of the 3’ end of U6 permitted intron turnover, indicating that RNA is not required in the loop region of the U6 ISL or in the bulge between the ISL and U2/U6 helix II (Supplemental Fig. S5A; see below). To refine the 5’ boundary of where RNA is required in the 3’ end of U6 for intron turnover, we tiled 8 nt blocks of DNA in 1 nt. steps 9, 10, and 11 nts from the 3’ end. These substitutions conferred diminishing consequences for intron turnover, with the impact of DNA substitutions on intron turnover dropping off markedly between nucleotides 96 and 97 (Fig. 5E, F, cf. 96-103D vs. 97-104D, lanes 9 and 10). Thus, the dependence of intron turnover and spliceosome disassembly on RNA at the 3’ end of U6 is confined to a short stretch of sixteen nucleotides, which must be RNA. Given evidence that small, 8-nt. DNA blocks can inhibit DEAH box ATPases (Semlow et al. 2016), these data are consistent with a model in which an RNA-dependent helicase, such as Prp43p, binds to single-stranded RNA at the very 3’ end of U6 and translocates upstream in a 2’-hydroxyl-dependent fashion to disrupt interactions, such as U2/U6 helix II.

This model is consistent with our analysis of Prp43p/Ntr1p activity on idealized unwinding substrates. Specifically, we found that a 10-nt ssRNA overhang, analogous to the ssRNA at the 3’ end of U6, was sufficient to support unwinding of a 24 bp duplex and that an 8-nt ssDNA substitution at the 3’ end of the overhang precluded unwinding and did so by precluding binding of Prp43p to the substrate (Fig. 5G; Fig. S5B), consistent with an interpretation in which the 105-112D substitution impedes recruitment of Prp43p to U6. Additionally, we found that an 8 nt DNA substitution at the proximal end or in the middle of a duplex permitted binding of Prp43p/Ntr1p to the substrate but reduced unwinding more than 5 fold (Fig. 5H; Fig. S5B), consistent with an interpretation in which the 97-104D substitution specifically impedes unwinding of U2/U6 helix II by Prp43p.

We also found that a DNA substitution at the distal end of the duplex did not impede unwinding (Fig. 5H, lanes 16-19), paralleling the failure of DNA substitutions downstream of 97-104D but within U2/U6 helix II to impair spliceosome disassembly. These results suggest that these non-inhibitory DNA blocks either i) impede translocation by Prp43p along a region that is not required for unwinding (because unwinding has already occurred) or ii) allow translocation of Prp43p, because unwinding has already occurred, and DNA only inhibits translocation during unwinding or while performing work beyond translocation. Thus, we cannot rule out at this time a model in which Prp43p translocates along U6 beyond U2/U6 helix II.

### The 3’ end of U6 interacts directly with the DEAH box ATPase Prp43p

To test explicitly whether the 3’ end of U6 may serve as a substrate for Prp43p, we assayed for a direct interaction between U6 and Prp43p by site-specific crosslinking. Although a recent *in vitro* study implicated Prp43p as interacting with pre-mRNA but not snRNAs (Fourmann et al. 2016), a previous study reported interactions between Prp43p and U6 using CRAC (UV-crosslinking and analysis of cDNA) *in vivo* (Bohnsack et al. 2009); nevertheless, these U6 sites, captured at steady-state, did not reveal an interaction at the 3’ end of U6 and instead provided evidence for interactions centered just downstream of the 5’ stem loop and on the 3’ side of the ISL, which is flanked by regions that spliceosome disassembly did not require to be RNA (Figure S5A). Thus, we performed site-specific UV crosslinking with U6 containing a ^32^P label upstream of 3 consecutive, photoactivatable 4-thio-U (s^4^U) residues incorporated into the 3’ end of U6 (Fig. 6A). We depleted U6 from Prp19p-HA tagged yeast extract, reconstituted with s^4^U-substituted U6 (*s4U_104-106 U6), assembled splicing reactions with a Cy5-labeled, *ACT1* pre-mRNA substrate (Supplemental Fig. S6B) and then, after the reactions were irradiated with UV light, enriched for spliceosome-bound U6 by immunoprecipitation with anti-HA antibodies.

**Figure 6.**
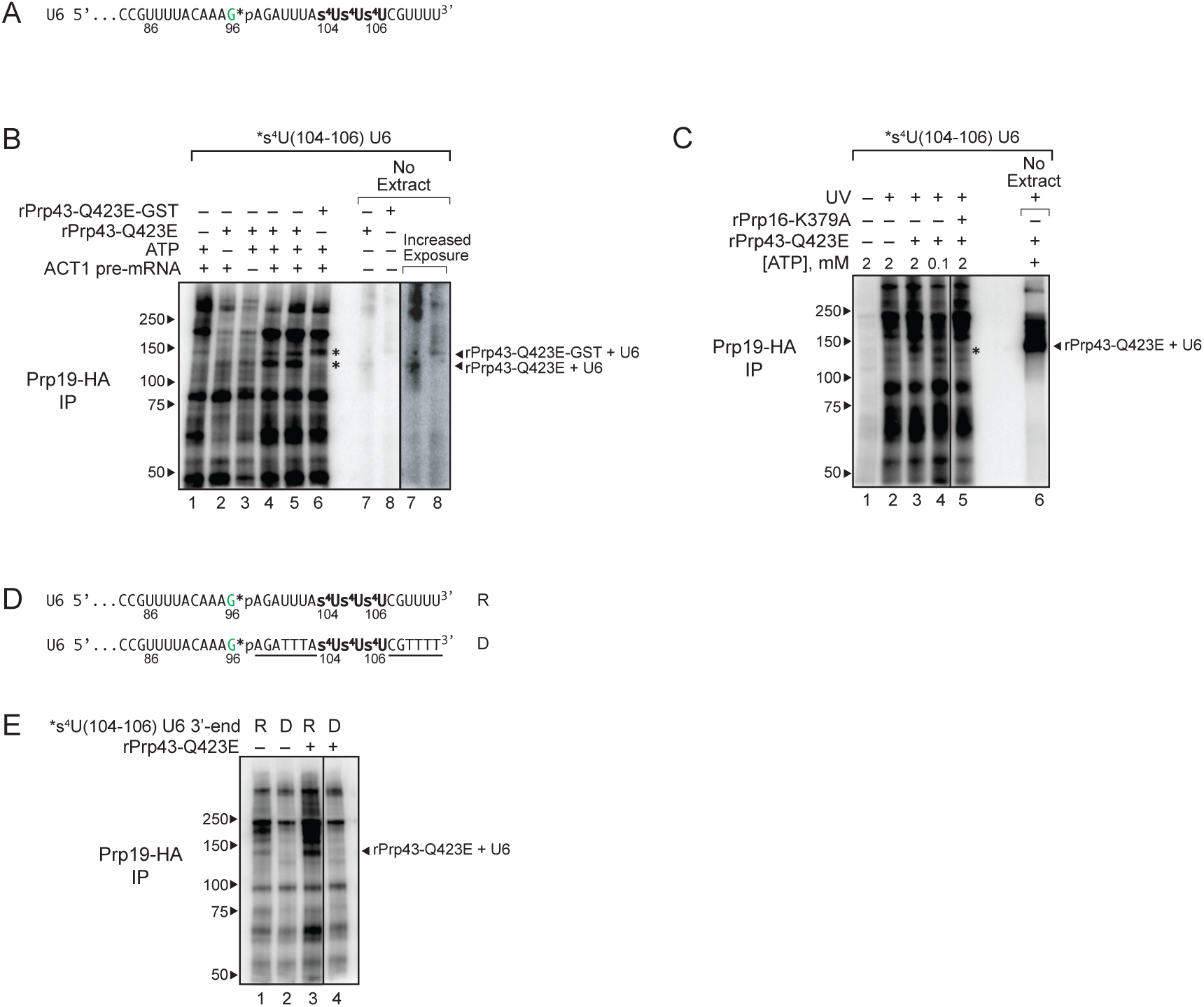
The 3’ end of U6 snRNA interacts directly with Prp43p at the disassembly stage of splicing. **A**. The 3’ end sequence of *s4U(104-106) U6, used for crosslinking, is shown with the location of 4-thio-U modifications (s^4^U) and the single ^32^P label (*p). **B,C**. U6 crosslinks to Prp43p in a splicing-dependent manner. Splicing of Cy5-*ACT1* pre-mRNA was performed under the indicated conditions in extracts (ySCC1) reconstituted with *s4U(104-106) U6 and supplemented where indicated with either rPrp43p-Q423E, GST-tagged rPrp43p-Q423E, or rPrp16p-K379A; splicing reactions were monitored by PAGE (Supplemental Fig. 6B); control reactions were performed in the absence of extract as indicated. After crosslinking, spliceosomes were immunoprecipitated (IP) under native conditions by HA-tagged Prp19p, and then crosslinks between U6 and spliceosomal proteins were analyzed by SDS-PAGE; control reactions were analyzed directly. Migration of protein size markers are indicated to the left, and the positions of U6 crosslinked to rPrp43-Q423E or rPrp43-Q423E-GST in the absence of extract are indicated to the right. Asterisk indicates migration of rPrp43-Q423E or rPrp43-Q423E-GST crosslinked to U6 in the absence of extract. See also Figs. S6 and S7. **D**. The 3’ end sequence of *s^4^U(104-106) U6, used for crosslinking, is shown with the location of 4-thio-U modifications (s^4^U), the single ^32^P label (*p), and DNA substitutions (underlined), where “R” indicates the U6 construct with RNA only at the 3’ end, and “D” indicates the U6 construct with the indicated DNA substitutions at the 3’ end. See also Figure S7. **E**. U6 crosslinks to Prp43p in an RNA-dependent manner. Splicing, crosslinking, and immunoprecipitation was executed and analyzed as in B and C except that the U6 variants shown in D were utilized.

In the absence of added rPrp43-Q423E, U6 forms a number of prominent crosslinks (Fig. 6B, lane 1). In the presence of rPrp43-Q423E, U6 formed an additional, prominent crosslink at ∼125 kDa, which corresponds to the molecular weight of Prp43p plus U6 snRNA (87.5 kDa + 37 kDa) (Fig. 6B, cf. lane 1 to lanes 4 and 5). Robust formation of this ∼125 kDa species, depended on both *ACT1* pre-mRNA and ATP (Fig. 6B, c.f. lanes 2 and 3 to 4 and 5), as well as UV and s^4^U (Fig. 6C; Supplementary Fig. S7). In the presence of rPrp43-Q423E tagged with GST, the ∼125 kDa crosslink disappeared and a new crosslink appeared at ∼150 kDa (Fig. 6B, lane 6), roughly equivalent to the increase in size expected for a GST tag (26 kDa). Supporting that these species reflect crosslinking of U6 to Prp43p, the mobilities of the ∼125 and ∼150 kDa crosslinks correlated to the mobilities of rPrp43-Q423E and rPrp43-Q423E-GST, respectively, crosslinked to U6 in the absence of yeast extract (Fig. 6B, cf. lanes 4 and 5 with lane 7, and lane 6 with lane 8).

Having identified U6 crosslinks in spliceosomes stalled prior to intron release, we next tested whether U6 crosslinked to Prp43p explicitly and, if so, whether such crosslinking occurred specifically at the 3’ end of U6. From the same crosslinking reactions described above, we pulled down His6-tagged Prp43p using Ni-NTA beads under denaturing conditions and then cleaved the 3’ end away from the remainder of U6 using RNase H targeted with an oligo complementary to nucleotides 48-93 (Supplemental Fig. S6C, D). In the presence of rPrp43-Q423E, the 3’ fragment of U6 was crosslinked in a species that migrated at ∼100 kDa, which corresponds roughly to the molecular weight of the fragment plus Prp43p (∼6.3 kDa + 87.5 kDa); this crosslink co-migrates with a crosslink between the 3’ fragment of U6 and rPrp43-Q423E formed in the absence of extract (Supplemental Fig. S6C). As with the ∼125 kDa species in the native spliceosome immunoprecipitates, substantial formation of this ∼100 kDa crosslink depended on both *ACT1* transcript and ATP, and the crosslink was shifted to a higher molecular weight of ∼125 kDa when extracts were instead supplemented with GST-tagged rPrp43-Q423E (Supplemental Fig. S6C). These crosslinking data confirm that U6 interacts directly with Prp43p and establish that U6 does so at its 3’ end.

To confirm that the interaction between U6 and Prp43p occurs at a late stage of splicing, we tested for dependence of crosslinking on earlier steps in splicing. When we allowed spliceosome assembly but impeded spliceosome activation by using a low concentration of ATP (0.1 µM, i.e. (Tarn et al. 1993)), U6 no longer crosslinked efficiently to rPrp43-Q423E, as compared to reactions with high concentrations of ATP (Fig. 6C, cf. lanes 3 and 4). When we permitted spliceosome activation but stalled splicing after the first chemical step of splicing with an ATPase-deficient Prp16p mutant (K379A) (Schneider et al. 2002), U6 again failed to crosslink efficiently to rPrp43-Q423E (Fig. 6C, cf. lane 3 and 5; Supplemental Fig. S7, cf. lanes 5 and 6). These data confirm that U6 interacts with Prp43p at a late stage in the splicing cycle.

Given that intron turnover requires that the 3’ end of U6 be RNA and not DNA (Fig. 5), to test whether Prp43p interacts with the 3’ end of U6 in a functional manner, we tested whether this interaction required RNA. Specifically, we assayed for crosslinking between Prp43p and U6 snRNA in which the last sixteen nucleotides (97-112) were replaced with DNA, with the exception of the three s^4^U-substituted nucleotides at 104-106 (Fig. 6D). This DNA substitution precluded crosslinking of Prp43p-Q423E to U6, indicating that the interaction of Prp43p with the 3’ end of U6 requires RNA (Fig. 6E, cf. lanes 3 and 4).

Because the region of U6 that must be RNA to promote intron turnover overlaps with U2/U6 helix II, we considered whether the sole function of the 3’ end of U6 at this stage is to enable Prp43p-mediated unwinding of U2/U6 helix II, which is present in the intron lariat spliceosome (ILS) (Wan et al. 2017; Zhang et al. 2019). If so, we would predict that disruption of U2/U6 helix II would bypass the requirement for Prp43p. We thus tested whether mutation of U6 nucleotides 93 to 102 to polyC (Fig. 7A) would suppress the intron turnover defect of rPrp43p-Q423E. This U6 variant did not suppress the intron turnover defect conferred by rPrp43p-Q423E (Fig. 7B), indicating that disruption of helix II is not sufficient for intron release and spliceosome disassembly. Together, our data implicate a model in which Prp43p disrupts interactions with U6 more broadly by pulling on the 3’ end of U6 to build up tension and thereby to disrupt interactions by acting at a distance (Figure 7C, see Discussion).

**Figure 7.**
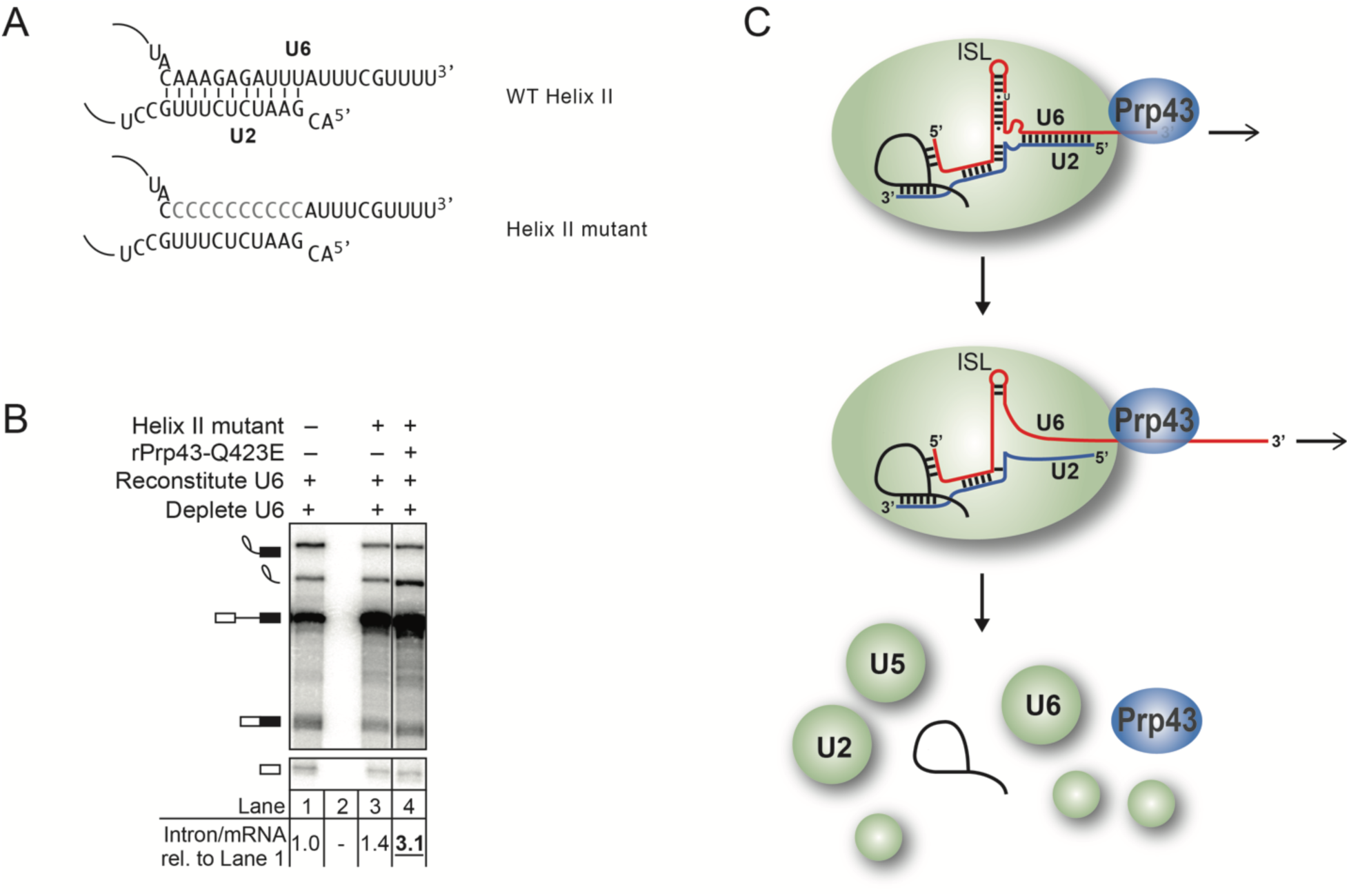
A model for Prp43 function on U6 involving disruption of structures beyond U2/U6 helix II on the 3’ end. **A**. U2/U6 helix II is depicted in the context of wild-type U6 or U6 with mutations that would disrupt base pairing to U2. **B**. Disruption of U2/U6 helix II does not bypass the requirement for Prp43p. *In vitro* splicing reactions were executed using radiolabeled *ACT1* pre-mRNA in extracts (yJPS866) reconstituted with wild-type or helix II mutant U6 (illustrated in panel A) and with or without rPrp43p-Q423E and then analyzed by denaturing PAGE. Quantitation of intron turnover is shown below the gel, wherein the ratio of excised intron to mRNA was calculated and normalized relative to the wild-type U6 control in lane 1; values with a substantial increase of 2-fold or greater over wild-type are indicated in bold, underlined. **C**. A mechanistic model for Prp43 function in splicing termination. See text for details.

## Discussion

Whereas identifying the functions of RNA-dependent SF2 ATPases in splicing has been relatively straightforward (Cordin and Beggs 2013), determining their targets and precise mechanisms has proven more challenging. Though we know the DEAH box ATPase Prp43 plays a critical role in terminating splicing, both to discard rejected suboptimal substrates and to release an excised optimal intron (Arenas and Abelson 1997; Martin et al. 2002; Koodathingal et al. 2010; Mayas et al. 2010; Pandit et al. 2006; Tsai et al. 2007), the mechanism of Prp43p action has been unclear (Fourmann et al. 2016; Wan et al. 2017; Zhang et al. 2019). We have discovered that the requirement for Prp43p in terminating splicing parallels a requirement for the 3’ end of U6 in the discard of a rejected lariat intermediate and in the release of an excised intron (Figs. 1, 2, and 3). The 3’ end of U6 is not required for the binding of the Prp43p-bound NTR complex to the spliceosome but rather a downstream step (Figs. 4). Consistent with a direct role for Prp43p at the 3’ end of U6 in splicing termination, Prp43p activity and splicing termination show the same nucleic acid preferences (Fig. 5). Indeed, Prp43p crosslinks to the 3’ end of U6 at a late stage of splicing in a manner that requires RNA nucleotides at the 3’ end (Fig. 6). Our data suggest a model in which Prp43 translocates along U6, leading to spliceosome disassembly through a pulling mechanism, as recently defined for two other DEAH box ATPases in splicing, Prp16p and Prp22p.

The first suggestion of a direct spliceosomal substrate for Prp43p derived from an *in vivo* crosslinking study, which revealed crosslinking of Prp43 to U6 snRNA (Bohnsack et al. 2009), in addition to rRNAs and snoRNAs, consistent with the additional role for Prp43p in ribosome biogenesis (Leeds et al. 2006; Combs et al. 2006; Lebaron et al. 2009). Subsequently, *in vitro* biochemical experiments suggested that Prp43p targets U2 snRNP for displacement from pre-mRNA and does so by interacting with pre-mRNA (Fourmann et al. 2016). While a recent cryo-EM structure of the yeast spliceosome at the disassembly stage did indicate proximity of Prp43p to the pre-mRNA intron, the structure revealed the closest proximity is to the 3’ end of U6, although direct interactions were not observed (Wan et al. 2017). A more recent cryo-EM structure of the human spliceosome also indicated proximity of Prp43p to the 3’ end of U6, although again no direct interactions were observed and Prp43p was no closer than 40Å from U6 (Zhang et al. 2019). Our data provide direct and functional evidence that Prp43p targets the 3’ end of U6 at the disassembly stage.

Evidence for two alternative models for Prp43p raises the question as to whether Prp43p functions in multiple capacities, as demonstrated for the RNA helicase DbpA, for example (Karginov and Uhlenbeck 2004). On the one hand, the model that Prp43 displaces U2 snRNP from pre-mRNA is founded on compelling data (Fourmann et al. 2016). First, the data supporting the model derive from experiments that leveraged a variant of Prp43p that is sufficient to disassemble the ILS in the absence of Ntr1p and Ntr2p, a variant that is constitutively activated by fusion of Prp43p to the G-patch domain of Ntr1p (Prp43p_Ntr1GP). Second, the model derives from direct and specific crosslinking from Prp43p_Ntr1GP to both pre-mRNA and to U2 snRNP proteins. Indeed, hPrp43p has been implicated as a component of the U2 snRNP – both in splicing (Will et al. 2002) and U7 biogenesis (Friend et al. 2007). Third, for all three spliceosomal intermediates tested, Prp43p_Ntr1GP-mediated disassembly consistently released the U2 snRNP. On the other hand, the model and supporting data are difficult to reconcile with several observations in the literature. First, the exclusive crosslinking of Prp43p_Ntr1GP to pre-mRNA does not align with the specific crosslinking of Prp43p to U6 *in vivo* (Bohnsack et al. 2009). Second, the data lead to a model in which Prp43p translocates 5’ to 3’ along the pre-mRNA to displace U2, but the DExH families of RNA-dependent ATPases translocate in the 3’ to 5’ direction (Gilman et al. 2017), and although earlier work provided some evidence for 5’ to 3’ translocation by Prp43p (Tanaka and Schwer 2006; Tanaka et al. 2007; Lebaron et al. 2009), more recent data using unwinding substrates with unstructured, single-stranded tails has established a strong preference for Prp43p in translocating in the 3’ to 5’ direction (He et al. 2017); additionally, the structure of the human spliceosome at the disassembly stage revealed proteins bound to the intron upstream of the U2-branchsite interaction, proteins that may preclude unwinding of this interaction by Prp43p (Zhang et al. 2019). Third, the crosslinking and disassembly data do not reflect the interaction and activity of Prp43p on spliceosomes poised for disassembly but rather the interaction and activity of Prp43p_Ntr1GP on spliceosomal intermediates that have not been identified as physiological targets for Prp43p either *in vivo* or in splicing extracts; for example, whereas the B^act^ complex is targeted by the Prp43p_Ntr1GP fusion for disassembly (Fourmann et al. 2016), this complex is not targeted by the NTR complex (Chen et al. 2013), presumably because SF3B1 sterically blocks Ntr2p binding (Rauhut et al. 2016; Yan et al. 2016); further, whereas the B complex is targeted by the Prp43p_Ntr1GP fusion for disassembly (Fourmann et al. 2016), and Prp43p and Ntr1p have been suggested to discard at this stage of spliceosome activation *in vivo* (Pandit et al. 2006), even before dissociation of the Lsm2-8 complex from the 3’ end of U6 (see below), the NTR complex is not sufficient to disassemble the B complex *in vitro* (Fourmann et al. 2016). Additional work will be required to resolve these discrepancies and to determine whether Prp43p functions exclusively through one mode or through multiple modes.

Our data define a model in which Prp43 releases a substrate by disassembling the spliceosome through translocation along U6, starting at the 3’ end (Fig. 7C). In particular, our crosslinking, binding, and DNA substitution data (Figs. 5,6, S5, and S6), indicate that Prp43p first binds the single-stranded tail of U6 and then translocates into U2/U6 helix II, disrupting this interaction. Unwinding of this helix likely contributes to spliceosome disassembly not just by initiating release of U2 from U6 but also by favoring formation of the 5’ stem loop of U2, which is mutually exclusive with U2/U6 helix I, a structure that includes the catalytic triad and juxtaposes the 5’ splice site and branch site sequences (Madhani and Guthrie 1992). Unwinding of U2/U6 helix II is not sufficient, however (Fig. 7A, B). In one model, limited translocation along the 3’ end of U6 would not only displace U2 but also build up tension that would destabilize upstream interactions, such as the catalytic U6 ISL and interactions between the ISL and Prp8p, a central splicing co-factor uniquely conserved from spliceosomes to self-splicing group II introns (Piccirilli and Staley 2016). Such dissociation at the heart of the spliceosome may then trigger disassembly of the remainder of the spliceosome, although both *in vivo* and sensitive *in vitro* assays have provided evidence for a role for the DExH box ATPase Brr2p as well (Small et al. 2006; c.f. Fourmann et al. 2013). A role for Prp43 in disrupting interactions by pulling on an RNA to build up tension and thereby disrupting interactions at a distance parallels recent evidence that the related DEAH box ATPases Prp16p and Prp22p pull on the substrate to dislodge regions of the substrate from the catalytic core of the spliceosome (Semlow et al. 2016). Consistent with this limited-pulling model, we have not yet identified regions of U6 upstream of U2/U6 helix II in which DNA substitutions inhibit Prp43p function (Fig. 5, S5). However, these DNA substitutions fall in primarily ssRNA regions. In an alternative model, Prp43p translocates more extensively along U6, performing disruptive work at regions yet to be defined. Consistent with this model, *in vivo* crosslinking identified interactions between Prp43p and upstream regions of U6 (Bohnsack et al. 2009). Defining the complete path of translocation and regions of disruptive work will require further studies.

Our functional data together with recent cryoEM structural data (Wan et al. 2017; Zhang et al. 2019) support a model by which Prp43p is targeted to its substrate and activated for spliceosome disassembly. In this model, Prp43p is localized in proximity to the 3’ end of U6 by binding to the NTC component Syf1p, and Prp43p is activated by binding of the conserved factor Ntr1p as well as Ntr2p in yeast to the spliceosome (Tsai et al. 2005; Boon et al. 2006; Tanaka et al. 2007), because binding appears to juxtapose the activating G-patch domain of Ntr1p with Prp43p; binding of Ntr1p and Ntr2p to the spliceosome may also derepress the NTR complex by reconfiguring interactions within the NTR complex (Fourmann et al. 2016; 2017). This model also rationalizes the action of Prp43p and the NTR complex at other stages of splicing, because Syf1p remains in proximity to the 3’ end of U6 in all spliceosomal complexes observed by cryoEM to-date (B^act^, B*, C, C*, P, and ILS) (Yan et al. 2019; Wan et al. 2019). Although the NTR complex functions as a general splicing terminator (Mayas et al. 2010; Koodathingal et al. 2010; Pandit et al. 2006; Chen et al. 2013), the NTR complex is restricted from acting at several stages of splicing. As noted, the binding and activation model rationalizes the inactivity of the NTR complex against the B^act^ complex and the C* and P complex, because Ntr2p binds to a pocket in Prp8 that is mutually exclusive with binding of SF3b1, in the B^act^ complex, and of Prp22p, in the C* and P complexes. Additionally (Wan et al. 2017; Zhang et al. 2019), the structure of the C complex rationalizes the inactivity of the NTR complex against this complex; Prp16p binds to the spliceosome in this complex in a manner that is mutually exclusive with the binding of Ntr1p and Ntr2p. The binding and or positioning of SF3b1p, Prp16p, and Prp22p likely signals productively engaged spliceosomes, whereas the prolonged absence of these factors likely signals unproductive, stalled spliceosomes, rendering them subject to NTR-mediated discard.

Interestingly, at the stage that Syf1p becomes stably bound to the spliceosome, as a component of the NTC complex, the Lsm2-8 complex dissociates from the 3’ end of U6 (Chan et al. 2003). Given the requirements for the 3’ end of U6 in spliceosome disassembly and the interaction of Prp43p with the 3’ end of U6, it is unlikely that Prp43p is active for disassembly of the spliceosome by translocation along U6 while the Lsm2-8 complex remains bound to U6. Thus, coincident with spliceosome activation in the transition from the B complex to B^act^ complex, both the docking site for Prp43p (Syf1p) and a target for Prp43p (3’ end of U6) become, in principle, available. In this manner, as the spliceosome becomes competent for catalysis, it also becomes primed for disassembly and termination, likely to ensure fidelity and recycling.

## Materials and Methods

### Strains

*In vitro* splicing was performed in whole-cell extracts of the following strains:

| Strain | Description | Genotype | Reference |
| --- | --- | --- | --- |
| yJPS860 | *PRP19-TAP* | *MATa his3Δ1 leu2Δ0 met15Δ0 ura3Δ0 prp19Δ::PRP19-TAP* | (Mayas et al. 2006) |
| ySCC1 | *PRP19-HA* | *MATa prc1 prb1 pep4 leu2 trp1 ura3 PRP19-HA* | (Chan et al. 2003) |
| yJPS1448 | *LSM3-V5* | *prp19::LEU2 [PRP19-HIS3] LSM3-V5::kanMX6* | This study |
| yJPS866 | *PRP24* | *MATα prp24::kanR met15Δ0 lys2Δ0 ura3Δ0 his3Δ1 leu2Δ0 [pSE362-PRP24]* | (Rader and Guthrie 2002) |
| yJPS867 | *PRP24-3HA* | *MATa prp24::kanR met15Δ0 lys2Δ0 ura3Δ0 his3Δ1 leu2Δ0 [pSE362-PRP24-3HA]* | (Rader and Guthrie 2002) |
| yJPS799 | *dbr1Δ* | *MATa his3Δ1 leu2Δ0 met15Δ0 ura3Δ0 dbr1Δ::kanMX4* | Open Biosystems |
| yJPS1405 | *PRP19-biotin* | *MATa his3Δ1 leu2Δ0 met15Δ0 ura3Δ0 prp19::HTB-KanR* | (Fica et al. 2013) |
| yJPS1481 | *prp8Δ* | *MATa his3Δ1 leu2Δ0 met15Δ0 ura3Δ0 prp8::LEU2 [PRP8-URA3]* | (Schellenberg et al. 2013) |

To make strains yJPS866 and yJPS867, strains GLS618 and GLS616 (Rader and Guthrie 2002) were transformed with either [pSE362-PRP24] or [pSE362-PRP24-3HA], respectively and the *PRP24-URA3* plasmid was shuffled out using 5-FOA.

To make strain yJPS1448, strain BY4741 (Open Biosystems) was transformed with the plasmid [*PRP19-URA3*], and then *PRP19* was replaced with *LEU2,* yielding the intermediate strain yJPS1276. Then, a V5 tag was integrated just downstream of *LSM3* using *KANMX6* as a marker, yielding the intermediate strain yJPS1431. Finally, the plasmid [*PRP19-URA3*] was replaced with the plasmid [*PRP19-HIS*] (bJPS2442), yielding yJPS1448.

To generate the yeast strain harboring HA-tagged Prp8p used in Figure S1, we transformed yJPS1481 with plasmid pJU204 (pSE362-PRP8-3HA; a gift from C. Guthrie) and shuffled out *PRP8-URA3* on 5-FOA and then streaked for single colonies on rich media.

### Preparation of recombinant proteins

Recombinant, dominant-negative His_6_-tagged rPrp43p-Q423E, was overexpressed from pJPS1384 (Mayas et al. 2010) in BL21 CodonPlus (DE3) RIL (Agilent) cells as described elsewhere (Semlow et al. 2016); the induced cells were lysed using a French press; recombinant protein was enriched via a Ni^+2^-nitroloacetic acid column (Ni-NTA; Qiagen), as described previously (Edwalds-Gilbert et al. 2000); and the eluted protein was fractionated on a 15-30% glycerol gradient, from which peak fractions were isolated, essentially as described elsewhere (Schneider et al. 2002; Semlow et al. 2016; Martin et al. 2002). Recombinant, dominant-negative rPrp22p-S635A, rPrp22p-K512A, and rPrp16p-K379A were expressed as described previously (Semlow et al. 2016).

To add an N-terminal GST tag to Prp43p-Q423E, the GST sequence from the pGEX-6P plasmid (GE Healthcare) was amplified using primers 5’-<u>GGCAGCAGCCATCATCATCATCATC ACAGCAGCGGCCTG</u>ATGTCCCCTATACTAGGTTATTGG-3’ and 5’-<u>TCTTCTTTTGGAACCCA TCATATGGCTGCCGCGCGGCAC</u>GGGCCCCTGGAACAGA-3’, which contain sequence that overlaps with the region between the His_6_-tag and N-terminus of Prp43p (underlined) from pJPS1384, with the remaining sequence overlapping with pGEX-6P. The product strands of this PCR amplification were subsequently used as megaprimers to perform overlap extension PCR cloning into pJPS1384 using Phusion DNA polymerase (Bryksin and Matsumura 2010). The insert was verified via Sanger sequencing, and the GST-tagged rPrp43p-Q423E protein was expressed and purified as described above.

Recombinant Prp43p and Ntr1p (1-120) proteins used for *in vitro* unwinding and EMSA experiments were expressed and purified as described previously (He et al. 2017).

### In vitro splicing, RNase H digestion, and U6 depletion/reconstitution

Wild-type *ACT1* pre-mRNA was transcribed from a HindIII-linearized pBST7ACTΔ6 plasmid (Mayas et al. 2006) using T7 RNA polymerase (Stevens and Abelson 2002) in the presence of ^32^P-UTP or, in the case of crosslinking experiments, Cy5-UTP. *UBC4* pre-mRNA having either a wild-type UAG or mutated UgG 3’ splice site were transcribed from PCR-generated DNA templates prepared by two rounds of PCR amplification. In the first round, pJPS2125 was amplified using primer 5’-GAACTAAGTGATCTAGAAAGG-3’ paired with either 5’-AACATGAAGTAGGTGGATCTCTAGTTCAATAGCAT-3’ for wild-type UAG or 5’-AACATGAA GTAGGTGGATCTC<u>C</u>AGTTCAATAGCAT-3’ for mutated U<u>g</u>G (underlined). The first round PCR products were then used as templates for the second round of amplification in which primer 5’-TAATACGACTCACTATAGGGAGAACTAAGTGATCT-3’ was paired with either the UAG or UgG primers listed above to introduce a T7 promoter for transcription (Mayas et al. 2010).

Yeast whole-cell extracts were prepared using the liquid nitrogen method as described elsewhere (Umen and Guthrie 1995), with modifications (Mayas et al. 2006). *In vitro* splicing reactions were performed under standard splicing conditions (Mayas et al. 2006) using 2 mM ATP (except where indicated) and 0.4-4 nM ^32^P-body-labeled *ACT1* or *UBC4* (having either a wild-type UAG or mutated UgG 3’ splice site) pre-mRNA substrate for 15-30 min at 20 °C. Where noted, splicing reactions were supplemented with recombinant, dominant-negative rPrp43p-Q423E, rPrp43p-Q423E-GST, rPrp22p-K512A, rPrp22p-S635A, or rPrp16p-K379A in buffer D (20 mM HEPES [pH 7.9], 0.2 mM EDTA, 50 mM KCl, 20% v/v glycerol, 0.5 mM DTT), at a final concentration of approximately 20-100 ng/µL (0.2-1 µM). Following *in vitro* splicing, RNA products were extracted using phenol/chloroform and separated on 6% (for *ACT1* substrate) or 15% (for *UBC4* substrates) denaturing polyacryamide gels.

For experiments testing the consequences of 3’ end cleavage of endogenous U6, a DNA oligo (or DNA-2’-O-methyl chimeric oligo) complementary to the targeted region was incubated at a concentration of approximately 10-20 µM with splicing reactions lacking ATP and substrate at 30 °C for 30 min (Fabrizio et al. 1989; Lapham et al. 1997). Following cleavage of U6 nucleotides via endogenous RNase H activity in the extract, an aliquot was removed to evaluate cleavage via northern blotting using a radiolabeled d1 oligo probe complementary to U6 nucleotides 28-54 (Small et al. 2006). Reactions were then supplemented with the appropriate amounts of ATP and splicing substrate and splicing was carried out as described above.

Depletion of endogenous U6 and reconstitution with T7-transcribed or synthetic wild-type U6, chimeric RNA-DNA U6 or s^4^U-substituted U6 was performed essentially as described (Fica et al. 2013; 2014). All observations are validated by redundant experiments and/or experiments that were reproduced at least twice.

### Glycerol gradients

Splicing reactions in a volume of 100 µL were assembled (following RNase H-directed cleavage of the 3’ end of endogenous U6, where indicated; see method above) in the presence of 2 nM radiolabeled *ACT1* or *UBC4* pre-mRNA and incubated at 20 °C for 30 min. Following incubation, 5 µL was removed and quenched as an input control, and an additional 5 µL was removed for evaluating cleavage of U6 via northern blot. The remaining reaction volume was then loaded onto hand-poured, 11 mL 15-40% glycerol gradients (20 mM HEPES [pH 7.9], 100 mM KCl, 2 mM MgCl_2_) and centrifuged at 37,000 rpm for 12-14 hours (Mayas et al. 2010). Aliquots of 440 µL were withdrawn from the top of the gradients, and RNA was extracted and fractionated on 6% (*ACT1* pre-mRNA) or 15% (*UBC4* pre-mRNA) denaturing polyacrylamide gels. For 15% gels, the gels were removed from plates and exposed wet to phosphorimager screens overnight in the cold room. See Data analysis below for description of gel imaging and analysis.

### Immunoprecipitations and affinity pull-downs

For immunoprecipitation of Prp43p-, Ntr1p-, Ntr2p-, and Prp24p-associated spliceosomes or snRNPs under native conditions (James et al. 2002), 25 µL splicing reactions were assembled (with or without RNase-H directed cleavage of U6; see method above) and incubated at 20 °C for 30 min. After removing a 5 µL aliquot as an input control, the remaining reaction volume was diluted 1:4 with IPP_150_ (10 mM Tris-HCl, pH 8, 150 mM NaCl, and 0.1% NP-40 substitute (Fluka)) and added to 5-7 µL of protein A sepharose beads (Sigma) in a 50% slurry from the vendor that had been washed in IPP_150_ and conjugated to ∼5-10 µg of affinity-purified or protein A-purified anti-Prp43p, anti-Ntr1p, or anti-Ntr2p antibodies (gifts from B. Schwer), and incubated with nutation for 1 hr at 4 °C. After incubation, beads were washed two times with 200 µL IPP_150_ and then two times with 500 µL of IPP_150_ for Prp43p, Ntr1p, and Ntr2p immunoprecipitation (IP) or just two times with 500 µL of IPP_150_ for all other IPs. RNA was extracted from the beads using phenol/chloroform.

For immunoprecipitation of spliceosomes or snRNPs from extract of the yeast strains *PRP19*-HA, *LSM3*-V5, or *PRP24*-3HA, 6-12 µg anti-HA antibodies (i.e., 12CA5, University of Chicago Monoclonal Antibody Facility) or 6 µg anti-V5 antibodies (R960-25, Invitrogen) per reaction were first conjugated to protein A sepharose beads in a 50% slurry that was first washed in IPP_150_. Immunoprecipitation was carried out as described above.

For affinity pull downs of His6-tagged rPrp43p-Q423E or His6-GST-tagged rPrp43-Q423E from UV-crosslinked splicing reactions under denaturing conditions, 5 µL of a 50% slurry of Ni-NTA beads were first washed in 20 volumes of wash buffer I (6 M guanidine•HCl, 50 mM [Tris pH 7.8], 300 mM NaCl, 0.1% NP-40 substitute, 10 mM imidazole, 5 mM betamercaptoethanol (BME))(Granneman et al. 2009). To the washed beads suspended in 50 µL of wash buffer I, 20 µL of UV-crosslinked splicing reaction was added and incubated for 3 hours with nutation at 4 °C. Following incubation, beads were washed twice with 250 µL of wash buffer I and then washed twice with 250 µL of wash buffer II (50 mM Tris [pH 7.8], 50 mM NaCl, 0.1% NP-40 substitute, 10 mM imidazole, and 5 mM BME) and treated with RNase H and analyzed by sodium dodecyl sulfate polyacrylamide gel electrophoresis (SDS-PAGE) as described below. RNA was extracted from the beads using phenol/chloroform.

### Synthesis of U6 for in vitro reconstitution and crosslinking

The 5’ piece of chimeric U6 snRNAs was constructed either by transcribing the first 96 nucleotides of U6 with T7 polymerase (Stevens and Abelson 2002) or by ligating the first 91 nucleotides of U6 in two sequential splint-mediated ligation steps from 3 pieces of synthetic RNA (Fica et al. 2014) or synthetic RNA substituted with blocks of DNA (Dharmacon). Synthetic oligonucleotides were first deprotected and gel purified prior to ligation. U6 oligonucleotides corresponding to U6 residues 97-112 or residues 92-112, made from 5’ phosphorylated synthetic RNA or synthetic RNA with various blocks of nucleotides substituted with DNA (Dharmacon or Integrated DNA Technologies), were then joined to U6 1-96 or U6 1-91, respectively, via a final splinted ligation. All ligation steps were performed with T4 DNA ligase (New England Biolabs) and DNA splints (Integrated DNA Technologies) with 20 nucleotides complementary to each side of the ligation junction in a 1:1:1 ratio of 5’ RNA:3’ RNA:splint. RNA pieces and splint were first annealed in the presence of 1X TEN buffer (10mM Tris-HCl, pH 7.5; 1mM EDTA; 66mM NaCl) by heating to 90 °C in a PCR block, and then slowing cooling to 18 °C at a rate of 1 °C per minute. T4 DNA ligase buffer, RNase inhibitor at 5% of the reaction volume (RNasin, Promega), and T4 DNA ligase (4,000 U per 50 µL reaction) were then added and incubated at 37 °C for 4-6 hours. Reactions were separated on a denaturing polyacrylamide gel (7M urea), ligation products were identified by UV-shadowing briefly at 254 nm, excised, eluted overnight at 4 °C in 1X TE (10mM Tris-HCl, pH 7.5; 1mM EDTA), and ethanol precipitated.

Note that for designing U6 snRNA with mutations in the region of U2/U6 helix II, we avoided polyA and polyG due to their tendency to form secondary interactions (i.e. base-stacking and quartets, respectively). For one DEAH-box helicase, MLE, a preference for polyU as well as poly(CU) has been observed (Prabu et al. 2015), so we chose not to use polyC as a comparison.

For crosslinking substrates, a synthetic oligonucleotide corresponding to U6 97-112, with 4-thio-U (s^4^U) substitutions at positions 104 to 106 (Dharmacon), was first deprotected and phenol-chloroform extracted and then 5’ end radiolabeled with [γ-^32^P] ATP (PerkinElmer, 6000 Ci/mmol) using T4 PNK (New England Biolab or Thermofisher) according to the manufacturer’s protocol. Following PNK treatment, the oligo was purified using a G-25 spin column, phenol-chloroform extracted, ethanol precipitated, and resuspended in 1X TE. The radiolabeled oligo was then ligated as described above to the previously assembled U6 1-96 in a ratio of 1.4:1:1.25 (U6 1-96:U6 97-112:splint), with the exception that ligation products were identified from a phosphorimage scan of the wet gel instead of UV-shadowing.

### In vitro unwinding and binding experiments

Substrates for unwinding and EMSA experiments were made by annealing RNA oligos (purchased from Sigma-Aldrich) to a 24mer DNA oligo containing a Quasar 670 (Cy5) fluorophore at the 3’ end (Biosearch Technologies) that was purified as described previously (He et al., 2017).

RNA oligos:

RNA-1: 5’GGCACCAACACAAAACACAUCUACACUCAACAAU

RNA-2: 5’GGCACCAACACAAAACACAUCUACAC[dT][dC][dA][dA][dC][dA][dA][dT]

RNA-3: 5’GGCACCAACACAAAACACAUCUACACUCAACAAUCACUUU

RNA-4: 5’ GGCACCAACACAAAAC[dA][dC][dA][dT][dC][dT][dA][dC]ACUCAACAAUCACUUU

RNA-5: 5’ GGCACCAA[dC][dA][dC][dA][dA][dA][dA][dC]ACAUCUACACUCAACAAUCACUUU

RNA-6: 5’[dG][dG][dC][dA][dC][dC][dA][dA]CACAAAACACAUCUACACUCAACAAUCACUUU

The sequence of the RNA oligos were designed as previously described (He et al., 2017) to avoid secondary structures.

DNA oligos:

Cy5 labelled oligo: 5’GTAGATGTGTTTTGTGTTGGTGCC-Cy5

Unlabeled oligo: 5’GTAGATGTGTTTTGTGTTGGTGCC

Capture oligo: 5’ GGCACCAACACAAAACACATCTAC

Unwinding reactions contained 4 nM substrate, 100 nM rPrp43p and 200 nM of its cofactor rNtr1p (1–120) in 40 mM Tris-HCl (pH 7.0), 0.5 mM MgCl2, and 2 mM DTT and in addition 80 nM unlabeled oligo to prevent reannealing of the Cy5 labeled oligo to the RNA oligo after unwinding. Unwinding was initiated by the addition of 1 mM ATP/MgCl_2_, and reactions were incubated on ice; alternatively, 1 mM ADP/MgCl_2_ was added as a control. Samples having a volume of 10 µL were removed at various time points and quenched with 5 µL of 3x loading buffer (5% Ficoll-type 400, 25% glycerol, 1.5X TBE, 20 mM EDTA, 0.5% SDS, 0.1% NP40 (Sigma 74385) and 160x excess of capture oligo) and analyzed on a native 10% polyacrylamide gel in 0.5X TBE (44.5 mM Tris-borate, 1 mM EDTA) containing 0.1 % SDS. Binding reactions (10 µL) for elecromobility shift assay (EMSA) reactions were assembled identically to unwinding reactions except that the 1 mM ATP/MgCl_2_ was omitted. Reactions were incubated on ice for 10 min, after which 5 µL of loading buffer (50% glycerol, 0.2% Triton X-100, 20 mM EDTA) was added, and reactions were analyzed on a 8% polyacrylamide gel in 0.25X TBE.

In Fig. 5G, RNA-1 and RNA-2 were annealed with the Cy5-labelled oligo to generate substrate-1 and substrate-2, respectively. In Fig. 5H, RNA-3 to RNA-6 were annealed with the Cy5-labelled oligo to generate substrate-3 to substrate-6, respectively.

### UV Crosslinking

Splicing reactions were reconstituted with s^4^U-substituted U6 snRNA (see above) at 15 fmol/5 µL reaction. Reactions were assembled with 2 nM *ACT1* pre-mRNA body-labeled with Cy5. Aliquots of each reaction were removed and analyzed on a splicing gel. The remainder of the reaction (∼30 µL) was then pipetted onto an aluminum block on ice and covered in parafilm and exposed to UV light (365 nm) at a distance of ∼3 cm for 20 minutes in a darkened room under an aluminum foil tent at 4 °C (Sontheimer 1994). Reactions were then subjected to immunoprecipitation under native conditions using anti-HA antibodies or to affinity pull-down under denaturing conditions using Ni-NTA beads. In the case of the denaturing pull down, after the final wash step (see method above) samples were digested with RNase H (Thermo Scientific) targeting U6 nucleotides 28-54 and 48-93 after incubation with 2.5 mM EDTA at 65 °C for 5 minutes to inactivate residual DNase I from the U6 depletion. All reactions were quenched with 6X SDS loading buffer and fractionated on an 8% tris-glycine SDS-PAGE gel along with a protein ladder (BioRad, Precision Plus Dual Color Standard). A photograph of the dried protein gel was overlaid with the scanned phosphorimage of the gel to estimate the molecular weights of bands on the phosphorimage.

### Primer extension analysis of UBC4 splicing reactions

Following slow-annealing to ∼500 fmol of ^32^P-radiolabeled primer oJPS239, RNA from splicing reactions (∼20 fmol pre-mRNA in 10.5 µL splicing reaction) was analyzed by primer extension (15 µL reaction volume total) using AMV reverse transcriptase (Promega) as previously described (Mayas et al. 2010). Products were analyzed on an 8% denaturing polyacrylamide sequencing gel.

### Data analysis

Gels were dried, except where indicated, and exposed to storage phosphorimager screens (Amersham Biosciences) for 24 to 48 hours and scanned on a Typhoon phosphorimager (Amersham Biosciences). For splicing gels of crosslinking reactions or gels of *in vitro* unwinding reactions and EMSA experiments, which used either Cy5-lableled splicing substrate or Cy5-lableled unwinding substrates, respectively, the gel was scanned wet within the two gels plates using the red laser on a Typhoon phosphorimager. Bands were quantified using TotalLab Quant software (version 12.2, TotalLab) using an automated rolling ball background subtraction algorithm.

## Acknowledgements

We thank B. Schwer for antibodies and C. Guthrie for strains. We thank D. Semlow for expressing and purifying recombinant Prp22p and Prp16p proteins. We thank Yi Zeng for help with data analysis and members of the Staley lab for comments on the manuscript. This work was supported by an NIH NRSA Individual Postdoctoral Fellowship (F32GM10192) awarded to RT and an NIH Research Project Grant (R01GM062264) awarded to JPS.

### Author contributions

R.T. and J.P.S. designed the overall study; R.T. performed all experiments except for the *in vitro* unwinding and binding experiments, which were performed by K.H.N; data analysis and interpretation was performed by R.T. and J.P.S., except for data analysis for unwinding experiments which was performed by K.H.N. and J.P.S. The manuscript was written, reviewed, and edited by R.T. and J.P.S., with contributions by K.H.N.

## Approval Statement

All authors have approved the current version of the manuscript and have approved its submission to *Genes & Development*.

## References

Achsel T, Brahms H, Kastner B, Bachi A, Wilm M, Luhrmann R. 1999. A doughnut-shaped heteromer of human Sm-like proteins binds to the 3’-end of U6 snRNA, thereby facilitating U4/U6 duplex formation in vitro. EMBO J 18: 5789–5802.

Appleby TC, Anderson R, Fedorova O, Pyle AM, Wang R, Liu X, Brendza KM, Somoza JR. 2011. Visualizing ATP-dependent RNA translocation by the NS3 helicase from HCV. J Mol Biol 405: 1139–1153.

Aravind L, Koonin EV. 1999. G-patch: a new conserved domain in eukaryotic RNA-processing proteins and type D retroviral polyproteins. Trends Biochem Sci 24: 342–344.

Arenas JE, Abelson JN. 1997. Prp43: An RNA helicase-like factor involved in spliceosome disassembly. Proc Natl Acad Sci USA 94: 11798–11802.

Bai R, Yan C, Wan R, Lei J, Shi Y. 2017. Structure of the Post-catalytic Spliceosome from Saccharomyces cerevisiae. Cell 171: 1589–1598.

Bertram K, Agafonov DE, Liu W-T, Dybkov O, Will CL, Hartmuth K, Urlaub H, Kastner B, Stark H, Lührmann R. 2017. Cryo-EM structure of a human spliceosome activated for step 2 of splicing. Nature 542: 318–323.

Bohnsack MT, Martin R, Granneman S, Ruprecht M, Schleiff E, Tollervey D. 2009. Prp43 Bound at Different Sites on the Pre-rRNA Performs Distinct Functions in Ribosome Synthesis. Molecular Cell 36: 583–592.

Boon KL, Auchynnikava T, Edwalds-Gilbert G, Barrass JD, Droop AP, Dez C, Beggs JD. 2006. Yeast ntr1/spp382 mediates prp43 function in postspliceosomes. Mol Cell Biol 26: 6016– 6023.

Bordonne R, Guthrie C. 1992. Human and human-yeast chimeric U6 snRNA genes identify structural elements required for expression in yeast. Nucleic Acids Res 20: 479–485.

Bryksin AV, Matsumura I. 2010. Overlap extension PCR cloning: a simple and reliable way to create recombinant plasmids. Biotechniques 48: 463–465.

Burgess SM, Guthrie C. 1993. A mechanism to enhance mRNA splicing fidelity: the RNA-dependent ATPase Prp16 governs usage of a discard pathway for aberrant lariat intermediates. Cell 73: 1377–1391.

Chan SP, Kao DI, Tsai WY, Cheng SC. 2003. The Prp19p-associated complex in spliceosome activation. Science 302: 279–282.

Chen H-C, Tseng C-K, Tsai R-T, Chung C-S, Cheng S-C. 2013. Link of NTR-mediated spliceosome disassembly with DEAH-box ATPases Prp2, Prp16, and Prp22. Mol Cell Biol 33: 514–525.

Chen MC, Tippana R, Demeshkina NA, Murat P, Balasubramanian S, Myong S, Ferré-D’Amaré AR. 2018. Structural basis of G-quadruplex unfolding by the DEAH/RHA helicase DHX36. Nature 558: 465–469.

Chen W, Shulha HP, Ashar-Patel A, Yan J, Green KM, Query CC, Rhind N, Weng Z, Moore MJ. 2014a. Endogenous U2·U5·U6 snRNA complexes in S. pombe are intron lariat spliceosomes. RNA 20: 308–320.

Chen Y-L, Capeyrou R, Humbert O, Mouffok S, Kadri YA, Lebaron S, Henras AK, Henry Y. 2014b. The telomerase inhibitor Gno1p/PINX1 activates the helicase Prp43p during ribosome biogenesis. Nucleic Acids Res 42: 7330–7345.

Combs DJ, Nagel RJ, Ares M, Stevens SW. 2006. Prp43p is a DEAH-box spliceosome disassembly factor essential for ribosome biogenesis. Mol Cell Biol 26: 523–534.

Company M, Arenas J, Abelson J. 1991. Requirement of the RNA helicase-like protein PRP22 for release of messenger RNA from spliceosomes. Nature 349: 487–493.

Cordin O, Beggs JD. 2013. RNA helicases in splicing. RNA Biol 10: 83–95.

Edwalds-Gilbert G, Kim DH, Kim SH, Tseng YH, Yu Y, Lin RJ. 2000. Dominant negative mutants of the yeast splicing factor Prp2 map to a putative cleft region in the helicase domain of DExD/H-box proteins. RNA 6: 1106–1119.

Faber ZJ, Chen X, Gedman AL, Boggs K, Cheng J, Ma J, Radtke I, Chao J-R, Walsh MP, Song G, et al. 2016. The genomic landscape of core-binding factor acute myeloid leukemias. Nat Genet 48: 1551–1556.

Fabrizio P, Dannenberg J, Dube P, Kastner B, Stark H, Urlaub H, Luhrmann R. 2009. The evolutionarily conserved core design of the catalytic activation step of the yeast spliceosome. Mol Cell 36: 593–608.

Fabrizio P, McPheeters DS, Abelson J. 1989. In vitro assembly of yeast U6 snRNP: a functional assay. Genes Dev 3: 2137–2150.

Fairman-Williams ME, Guenther U-P, Jankowsky E. 2010. SF1 and SF2 helicases: family matters. Curr Opin Struct Biol 20: 313–324.

Fica SM, Mefford MA, Piccirilli JA, Staley JP. 2014. Evidence for a group II intron-like catalytic triplex in the spliceosome. Nat Struct Mol Biol 21: 464–471.

Fica SM, Nagai K. 2017. Cryo-electron microscopy snapshots of the spliceosome: structural insights into a dynamic ribonucleoprotein machine. Nat Struct Mol Biol 24: 791–799.

Fica SM, Oubridge C, Galej WP, Wilkinson ME, Bai X-C, Newman AJ, Nagai K. 2017. Structure of a spliceosome remodelled for exon ligation. Nature 542: 377–380.

Fica SM, Tuttle N, Novak T, Li N-S, Lu J, Koodathingal P, Dai Q, Staley JP, Piccirilli JA. 2013. RNA catalyses nuclear pre-mRNA splicing. Nature 503: 229–234.

Fourmann J-B, Dybkov O, Agafonov DE, Tauchert MJ, Urlaub H, Ficner R, Fabrizio P, Lührmann R. 2016. The target of the DEAH-box NTP triphosphatase Prp43 in Saccharomyces cerevisiae spliceosomes is the U2 snRNP-intron interaction. eLife 5: e15564.

Fourmann J-B, Tauchert MJ, Ficner R, Fabrizio P, Lührmann R. 2017. Regulation of Prp43-mediated disassembly of spliceosomes by its cofactors Ntr1 and Ntr2. Nucleic Acids Res 45: 4068–4080.

Fourmann JB, Schmitzova J, Christian H, Urlaub H, Ficner R, Boon KL, Fabrizio P, Luhrmann R. 2013. Dissection of the factor requirements for spliceosome disassembly and the elucidation of its dissociation products using a purified splicing system. Genes Dev 27: 413– 428.

Friend K, Lovejoy AF, Steitz JA. 2007. U2 snRNP binds intronless histone pre-mRNAs to facilitate U7-snRNP-dependent 3’ end formation. Mol Cell 28: 240–252.

Fu X-D, Ares M. 2014. Context-dependent control of alternative splicing by RNA-binding proteins. Nat Rev Genet 15: 689–701.

Galej WP, Wilkinson ME, Fica SM, Oubridge C, Newman AJ, Nagai K. 2016. Cryo-EM structure of the spliceosome immediately after branching. Nature 537: 197–201.

Gilman B, Tijerina P, Russell R. 2017. Distinct RNA-unwinding mechanisms of DEAD-box and DEAH-box RNA helicase proteins in remodeling structured RNAs and RNPs. Biochem Soc Trans 45: 1313–1321.

Gottschalk A, Neubauer G, Banroques J, Mann M, Luhrmann R, Fabrizio P. 1999. Identification by mass spectrometry and functional analysis of novel proteins of the yeast [U4/U6.U5] tri-snRNP. EMBO J 18: 4535–4548.

Granneman S, Kudla G, Petfalski E, Tollervey D. 2009. Identification of protein binding sites on U3 snoRNA and pre-rRNA by UV cross-linking and high-throughput analysis of cDNAs. Proc Natl Acad Sci USA 106: 9613–9618.

Gu M, Rice CM. 2010. Three conformational snapshots of the hepatitis C virus NS3 helicase reveal a ratchet translocation mechanism. Proc Natl Acad Sci USA 107: 521–528.

Hamann F, Enders M, Ficner R. 2019. Structural basis for RNA translocation by DEAH-box ATPases. Nucleic Acids Res 3: 4349–4362.

Hang J, Wan R, Yan C, Shi Y. 2015. Structural basis of pre-mRNA splicing. Science 349: 1191–1198.

He Y, Andersen GR, Nielsen KH. 2010. Structural basis for the function of DEAH helicases. EMBO Rep 11: 180–186.

He Y, Staley JP, Andersen GR, Nielsen KH. 2017. Structure of the DEAH/RHA ATPase Prp43p bound to RNA implicates a pair of hairpins and motif Va in translocation along RNA. RNA 23: 1110–1124.

Heininger AU, Hackert P, Andreou AZ, Boon K-L, Memet I, Prior M, Clancy A, Schmidt B, Urlaub H, Schleiff E, et al. 2016. Protein cofactor competition regulates the action of a multifunctional RNA helicase in different pathways. RNA Biol 13: 320–330.

Hilliker AK, Staley JP. 2004. Multiple functions for the invariant AGC triad of U6 snRNA. RNA 10: 921–928.

James S-A, Turner W, Schwer B. 2002. How Slu7 and Prp18 cooperate in the second step of yeast pre-mRNA splicing. RNA 8: 1068–1077.

Jandrositz A, Guthrie C. 1995. Evidence for a Prp24 binding site in U6 snRNA and in a putative intermediate in the annealing of U6 and U4 snRNAs. EMBO J 14: 820–832.

Jankowsky E. 2011. RNA helicases at work: binding and rearranging. Trends Biochem Sci 36: 19–29.

Kannan R, Hartnett S, Voelker RB, Berglund JA, Staley JP, Baumann P. 2013. Intronic sequence elements impede exon ligation and trigger a discard pathway that yields functional telomerase RNA in fission yeast. Genes Dev 27: 627–638.

Kannan R, Helston RM, Dannebaum RO, Baumann P. 2015. Diverse mechanisms for spliceosome-mediated 3’ end processing of telomerase RNA. Nat Commun 6: 6104.

Karginov FV, Uhlenbeck OC. 2004. Interaction of Escherichia coli DbpA with 23S rRNA in different functional states of the enzyme. Nucleic Acids Res 32: 3028–3032.

Kim SH, Smith J, Claude A, Lin RJ. 1992. The purified yeast pre-mRNA splicing factor PRP2 is an RNA-dependent NTPase. EMBO J 11: 2319–2326.

Koodathingal P, Novak T, Piccirilli JA, Staley JP. 2010. The DEAH box ATPases Prp16 and Prp43 cooperate to proofread 5’ splice site cleavage during pre-mRNA splicing. Mol Cell 39: 385–395.

Krishnan R, Blanco MR, Kahlscheuer ML, Abelson J, Guthrie C, Walter NG. 2013. Biased Brownian ratcheting leads to pre-mRNA remodeling and capture prior to first-step splicing. Nat Struct Mol Biol 20: 1450–1457.

Lapham J, Yu YT, Shu MD, Steitz JA, Crothers DM. 1997. The position of site-directed cleavage of RNA using RNase H and 2’-O-methyl oligonucleotides is dependent on the enzyme source. RNA 3: 950–951.

Lebaron S, Froment C, Fromont-Racine M, Rain J-C, Monsarrat B, Caizergues-Ferrer M, Henry Y. 2005. The splicing ATPase prp43p is a component of multiple preribosomal particles. Mol Cell Biol 25: 9269–9282.

Lebaron S, Papin C, Capeyrou R, Chen Y-L, Froment C, Monsarrat B, Caizergues-Ferrer M, Grigoriev M, Henry Y. 2009. The ATPase and helicase activities of Prp43p are stimulated by the G-patch protein Pfa1p during yeast ribosome biogenesis. EMBO J 28: 3808–3819.

Leeds NB, Small EC, Hiley SL, Hughes TR, Staley JP. 2006. The splicing factor Prp43p, a DEAH box ATPase, functions in ribosome biogenesis. Mol Cell Biol 26: 513–522.

Licht K, Medenbach J, Luhrmann R, Kambach C, Bindereif A. 2008. 3’-cyclic phosphorylation of U6 snRNA leads to recruitment of recycling factor p110 through LSm proteins. RNA 14: 1532–1538.

Liu HL, Cheng SC. 2012. The Interaction of Prp2 with a Defined Region of the Intron Is Required for the First Splicing Reaction. Mol Cell Biol 32: 5056–5066

Liu S, Li X, Zhang L, Jiang J, Hill RC, Cui Y, Hansen KC, Zhou ZH, Zhao R. 2017. Structure of the yeast spliceosomal postcatalytic P complex. Science 358: 1278–1283.

Luo D, Xu T, Watson RP, Scherer-Becker D, Sampath A, Jahnke W, Yeong SS, Wang CH, Lim SP, Strongin A, et al. 2008. Insights into RNA unwinding and ATP hydrolysis by the flavivirus NS3 protein. EMBO J 27: 3209–3219.

Madhani HD, Guthrie C. 1992. A novel base-pairing interaction between U2 and U6 snRNAs suggests a mechanism for the catalytic activation of the spliceosome. Cell 71: 803–817.

Madhani HD, Guthrie C. 1994. Dynamic RNA-RNA interactions in the spliceosome. Annu Rev Genet 28: 1–26.

Martin A, Schneider S, Schwer B. 2002. Prp43 is an essential RNA-dependent ATPase required for release of lariat-intron from the spliceosome. J Biol Chem 277: 17743–17750.

Mayas RM, Maita H, Semlow DR, Staley JP. 2010. Spliceosome discards intermediates via the DEAH box ATPase Prp43p. Proc Natl Acad Sci USA 107: 10020–10025.

Mayas RM, Maita H, Staley JP. 2006. Exon ligation is proofread by the DExD/H-box ATPase Prp22p. Nat Struct Mol Biol 13: 482–490.

Montemayor EJ, Curran EC, Liao HH, Andrews KL, Treba CN, Butcher SE, Brow DA. 2014. Core structure of the U6 small nuclear ribonucleoprotein at 1.7-Å resolution. Nat Struct Mol Biol 21: 544–551.

Montemayor EJ, Didychuk AL, Yake AD, Sidhu GK, Brow DA, Butcher SE. 2018. Architecture of the U6 snRNP reveals specific recognition of 3’-end processed U6 snRNA. Nat Commun 9: 1–11.

Ohrt T, Odenwälder P, Dannenberg J, Prior M, Warkocki Z, Schmitzová J, Karaduman R, Gregor I, Enderlein J, Fabrizio P, et al. 2013. Molecular dissection of step 2 catalysis of yeast pre-mRNA splicing investigated in a purified system. RNA 19: 1–14.

Ozgur S, Buchwald G, Falk S, Chakrabarti S, Prabu JR, Conti E. 2015. The conformational plasticity of eukaryotic RNA-dependent ATPases. FEBS J 282: 850–863.

Pandit S, Lynn B, Rymond BC. 2006. Inhibition of a spliceosome turnover pathway suppresses splicing defects. Proc Natl Acad Sci USA 103: 13700–13705.

Pannone BK, Kim SD, Noe DA, Wolin SL. 2001. Multiple functional interactions between components of the Lsm2-Lsm8 complex, U6 snRNA, and the yeast La protein. Genetics 158: 187–196.

Piccirilli JA, Staley JP. 2016. Reverse transcriptases lend a hand in splicing catalysis. Nat Struct Mol Biol 23: 507–509.

Prabu JR, Müller M, Thomae AW, Schüssler S, Bonneau F, Becker PB, Conti E. 2015. Structure of the RNA Helicase MLE Reveals the Molecular Mechanisms for Uridine Specificity and RNA-ATP Coupling. Mol Cell 60: 487–499.

Pyle AM. 2008. Translocation and unwinding mechanisms of RNA and DNA helicases. Annu Rev Biophys 37: 317–336.

Qi X, Rand DP, Podlevsky JD, Li Y, Mosig A, Stadler PF, Chen JJ-L. 2015. Prevalent and distinct spliceosomal 3’-end processing mechanisms for fungal telomerase RNA. Nat Commun 6: 6105–6113.

Rader SD, Guthrie C. 2002. A conserved Lsm-interaction motif in Prp24 required for efficient U4/U6 di-snRNP formation. RNA 8: 1378–1392.

Raghunathan PL, Guthrie C. 1998. RNA unwinding in U4/U6 snRNPs requires ATP hydrolysis and the DEIH-box splicing factor Brr2. Curr Biol 8: 847–855.

Rauhut R, Fabrizio P, Dybkov O, Hartmuth K, Pena V, Chari A, Kumar V, Lee C-T, Urlaub H, Kastner B, et al. 2016. Molecular architecture of the Saccharomyces cerevisiae activated spliceosome. Science 353: 1399–1405.

Robert-Paganin J, Réty S, Leulliot N. 2015. Regulation of DEAH/RHA helicases by G-patch proteins. Biomed Res Int 2015: 931857–9.

Ryan DE, Stevens SW, Abelson J. 2002. The 5“ and 3” domains of yeast U6 snRNA: Lsm proteins facilitate binding of Prp24 protein to the U6 telestem region. RNA 8: 1011–1033.

Schellenberg MJ, Wu T, Ritchie DB, Fica S, Staley JP, Atta KA, LaPointe P, MacMillan AM. 2013. A conformational switch in PRP8 mediates metal ion coordination that promotes pre-mRNA exon ligation. Nat Struct Mol Biol 20: 728–734.

Schneider S, Hotz H-R, Schwer B. 2002. Characterization of dominant-negative mutants of the DEAH-box splicing factors Prp22 and Prp16. J Biol Chem 277: 15452–15458.

Schwer B, Gross CH. 1998. Prp22, a DExH-box RNA helicase, plays two distinct roles in yeast pre-mRNA splicing. EMBO J 17: 2086–2094.

Schwer B, Guthrie C. 1992. A conformational rearrangement in the spliceosome is dependent on PRP16 and ATP hydrolysis. EMBO J 11: 5033–5039.

Schwer B, Guthrie C. 1991. PRP16 is an RNA-dependent ATPase that interacts transiently with the spliceosome. Nature 349: 494–499.

Schwer B, Meszaros T. 2000. RNA helicase dynamics in pre-mRNA splicing. EMBO J 19: 6582–6591.

Scotti MM, Swanson MS. 2016. RNA mis-splicing in disease. Nat Rev Genet 17: 19–32.

Semlow DR, Blanco MR, Walter NG, Staley JP. 2016. Spliceosomal DEAH-Box ATPasesc Remodel Pre-mRNA to Activate Alternative Splice Sites. Cell 164: 985–998.

Semlow DR, Staley JP. 2012. Staying on message: ensuring fidelity in pre-mRNA splicing. Trends Biochem Sci 37: 263–273.

Shannon KW, Guthrie C. 1991. Suppressors of a U4 snRNA mutation define a novel U6 snRNP protein with RNA-binding motifs. Genes Dev 5: 773–785.

Small EC, Leggett SR, Winans AA, Staley JP. 2006. The EF-G-like GTPase Snu114p regulates spliceosome dynamics mediated by Brr2p, a DExD/H box ATPase. Mol Cell 23: 389–399.

Sontheimer EJ. 1994. Site-specific RNA crosslinking with 4-thiouridine. Mol Biol Rep 20: 35–44.

Staley JP, Guthrie C. 1999. An RNA switch at the 5’ splice site requires ATP and the DEAD box protein Prp28p. Mol Cell 3: 55–64.

Staley JP, Guthrie C. 1998. Mechanical devices of the spliceosome: motors, clocks, springs, and things. Cell 92: 315–326.

Stevens SW, Abelson J. 1999. Purification of the yeast U4/U6.U5 small nuclear ribonucleoprotein particle and identification of its proteins. Proc Natl Acad Sci USA 96: 7226–7231.

Stevens SW, Abelson J. 2002. Yeast pre-mRNA splicing: methods, mechanisms, and machinery. Methods Enzymol 351: 200–220.

Su Y-L, Chen H-C, Tsai R-T, Lin P-C, Cheng S-C. 2018. Cwc23 is a component of the NTR complex and functions to stabilize Ntr1 and facilitate disassembly of spliceosome intermediates. Nucleic Acids Res 46: 3764–3773.

Sun JS, Manley JL. 1995. A novel U2-U6 snRNA structure is necessary for mammalian mRNA splicing. Genes Dev 9: 843–854.

Tanaka N, Aronova A, Schwer B. 2007. Ntr1 activates the Prp43 helicase to trigger release of lariat-intron from the spliceosome. Genes Dev 21: 2312–2325.

Tanaka N, Schwer B. 2006. Mutations in PRP43 that uncouple RNA-dependent NTPase activity and pre-mRNA splicing function. Biochemistry 45: 6510–6521.

Tarn WY, Lee KR, Cheng SC. 1993. Yeast precursor mRNA processing protein PRP19 associates with the spliceosome concomitant with or just after dissociation of U4 small nuclear RNA. Proc Natl Acad Sci USA 90: 10821–10825.

Tauchert MJ, Fourmann J-B, Lührmann R, Ficner R. 2017. Structural insights into the mechanism of the DEAH-box RNA helicase Prp43. eLife 6: 1–25.

Tsai RT, Fu RH, Yeh FL, Tseng CK, Lin YC, Huang YH, Cheng SC. 2005. Spliceosome disassembly catalyzed by Prp43 and its associated components Ntr1 and Ntr2. Genes Dev 19: 2991–3003.

Tsai RT, Tseng CK, Lee PJ, Chen HC, Fu RH, Chang KJ, Yeh FL, Cheng SC. 2007. Dynamic interactions of Ntr1-Ntr2 with Prp43 and with U5 govern the recruitment of Prp43 to mediate spliceosome disassembly. Mol Cell Biol 27: 8027–8037.

Umen JG, Guthrie C. 1995. A novel role for a U5 snRNP protein in 3’ splice site selection. Genes Dev 9: 855–868.

Vidal VP, Verdone L, Mayes AE, Beggs JD. 1999. Characterization of U6 snRNA-protein interactions. RNA 5: 1470–1481.

Wagner JD, Jankowsky E, Company M, Pyle AM, Abelson JN. 1998. The DEAH-box protein PRP22 is an ATPase that mediates ATP-dependent mRNA release from the spliceosome and unwinds RNA duplexes. EMBO J 17: 2926–2937.

Wan R, Bai R, Yan C, Lei J, Shi Y. 2019. Structures of the Catalytically Activated Yeast Spliceosome Reveal the Mechanism of Branching. Cell 177: 339–351.

Wan R, Yan C, Bai R, Huang G, Shi Y. 2016. Structure of a yeast catalytic step I spliceosome at 3.4 Å resolution. Science 353: 895–904.

Wan R, Yan C, Bai R, Lei J, Shi Y. 2017. Structure of an Intron Lariat Spliceosome from Saccharomyces cerevisiae. Cell 171: 120–132.

Warkocki Z, Schneider C, Mozaffari-Jovin S, Schmitzová J, Höbartner C, Fabrizio P, Lührmann R. 2015. The G-patch protein Spp2 couples the spliceosome-stimulated ATPase activity of the DEAH-box protein Prp2 to catalytic activation of the spliceosome. Genes Dev 29: 94– 107.

Wilkinson ME, Fica SM, Galej WP, Norman CM, Newman AJ, Nagai K. 2017. Postcatalytic spliceosome structure reveals mechanism of 3’-splice site selection. Science 358: 1283– 1288.

Will CL, Lührmann R. 2011. Spliceosome structure and function. Cold Spring Harb Perspect Biol 3: 1–23.

Will CL, Urlaub H, Achsel T, Gentzel M, Wilm M, Lührmann R. 2002. Characterization of novel SF3b and 17S U2 snRNP proteins, including a human Prp5p homologue and an SF3b DEAD-box protein. EMBO J 21: 4978–4988.

Wlodaver AM, Staley JP. 2014. The DExD/H-box ATPase Prp2p destabilizes and proofreads the catalytic RNA core of the spliceosome. RNA 20: 1–13.

Wolff T, Bindereif A. 1995. Mutational analysis of human U6 RNA: stabilizing the intramolecular helix blocks the spliceosomal assembly pathway. Biochim Biophys Acta 1263: 39–44.

Wolin SL, Cedervall T. 2002. The La protein. Annu Rev Biochem 71: 375–403.

Yan C, Wan R, Bai R, Huang G, Shi Y. 2016. Structure of a yeast activated spliceosome at 3.5 Å resolution. Science 353: 904–911.

Yan C, Wan R, Bai R, Huang G, Shi Y. 2017. Structure of a yeast step II catalytically activated spliceosome. Science 355: 149–155.

Yan C, Wan R, Shi Y. 2019. Molecular Mechanisms of pre-mRNA Splicing through Structural Biology of the Spliceosome. Cold Spring Harb Perspect Biol 11: a032409.

Zhang X, Zhan X, Yan C, Zhang W, Liu D, Lei J, Shi Y. 2019. Structures of the human spliceosomes before and after release of the ligated exon. Cell Res 29: 274–285.

Zhou L, Hang J, Zhou Y, Wan R, Lu G, Yin P, Yan C, Shi Y. 2014. Crystal structures of the Lsm complex bound to the 3’ end sequence of U6 small nuclear RNA. Nature 506: 116–120.

Zuker M. 2003. Mfold web server for nucleic acid folding and hybridization prediction. Nucleic Acids Res 31: 3406–3415.

