## Supplemental Files for "Termination of pre-mRNA splicing requires that the ATPase and RNA unwindase Prp43 acts on the catalytic snRNA U6"

Toroney_FigS1

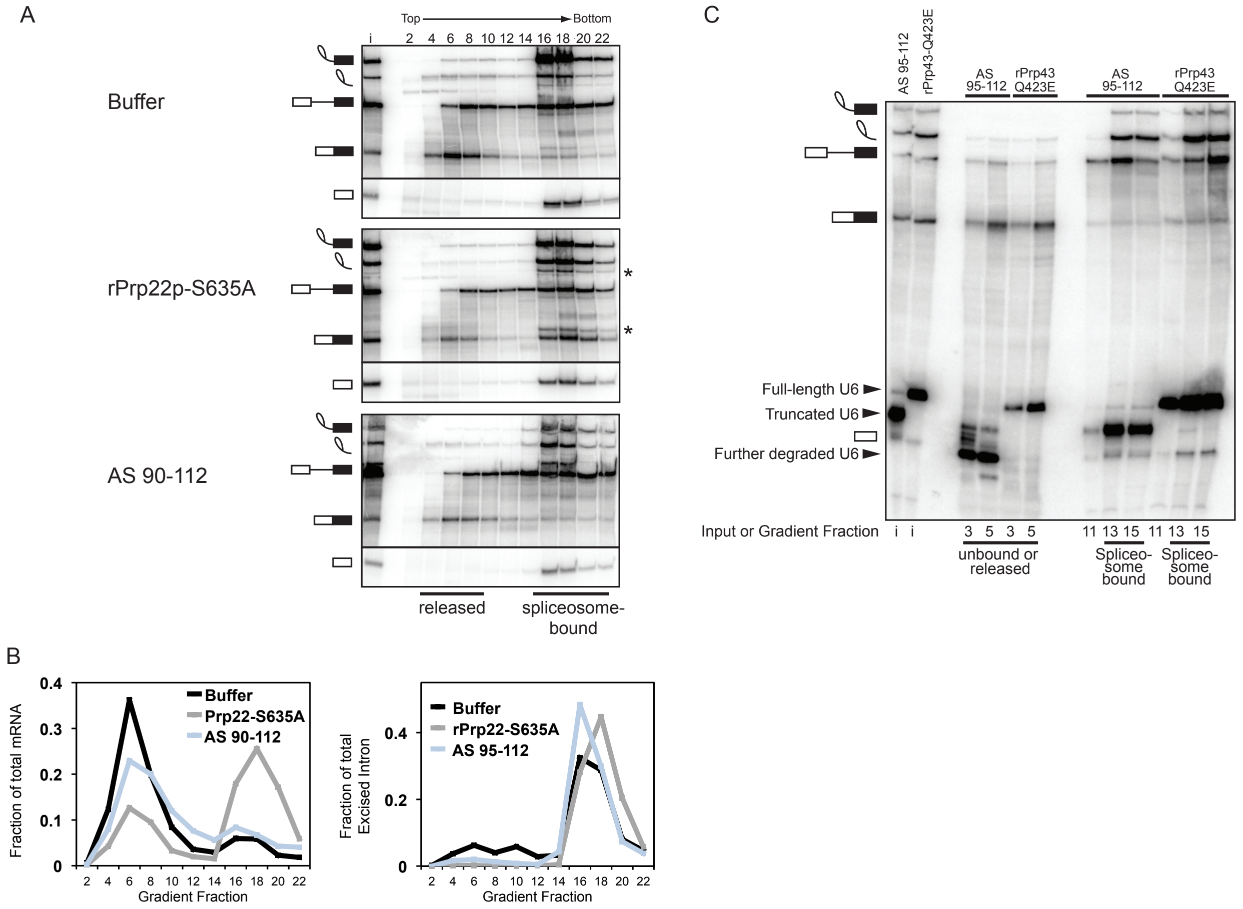

**<u>Figure S1</u>.** The 3' end of U6 is required for release of excised intron. **A**. Radiolabeled *ACT1* pre-mRNA was spliced in wild-type *DBR1*, Prp8-HA-tagged yeast extract (see Methods) that was subjected to RNase H cleavage of U6 directed by DNA oligo AS 90-112 (bottom panel) or that was supplemented with either buffer (top panel) or mutated rPrp22p-S635A (middle panel) (Schwer and Meszaros 2000). Splicing reactions were fractionated on a glycerol gradient; input (i) and fraction numbers are indicated above the top panel. Fractions containing released or spliceosome-bound splicing species are highlighted below the bottom panel. The asterisks indicate splicing at an upstream cryptic 3’ splice site, resulting from the proofreading defect of the Prp22p mutation (Mayas et al. 2006). **B**. Quantitation of mRNA (left) or excised intron (right) in each gradient fraction shown in **A**. Data are normalized as a fraction of total mRNA or total excised intron within each gradient. **C**. Truncated U6 snRNA assembles into spliceosomes that subsequently accumulate excised intron. Northern blot of fractions from gradients shown in Figure 2A, panels 2 and 4 (*dbr1Δ* [yJPS799] extracts). Even fractions are shown in Figure 2A; select, odd fractions are shown here and were run side-by-side on a separate denaturing PAGE gel along with inputs (i) and analyzed with a probe complementary to nucleotides 28-54 of U6. Position of full-length U6 as well as truncated U6, due to digestion by RNase H directed by AS 95-112, are indicated. Both full-length and truncated U6 comigrate with spliceosomes in the deep fractions (11-15), most unambiguously identified by lariat intermediates, indicating that truncated U6 is incorporated into spliceosomes and remains bound when release of the excised intron is blocked. Note that in shallow fractions containing RNAs not bound to the spliceosome (3 and 5) truncated U6 has degraded further, comigrating with and obscuring the 5' exon.

Toroney_FigS2

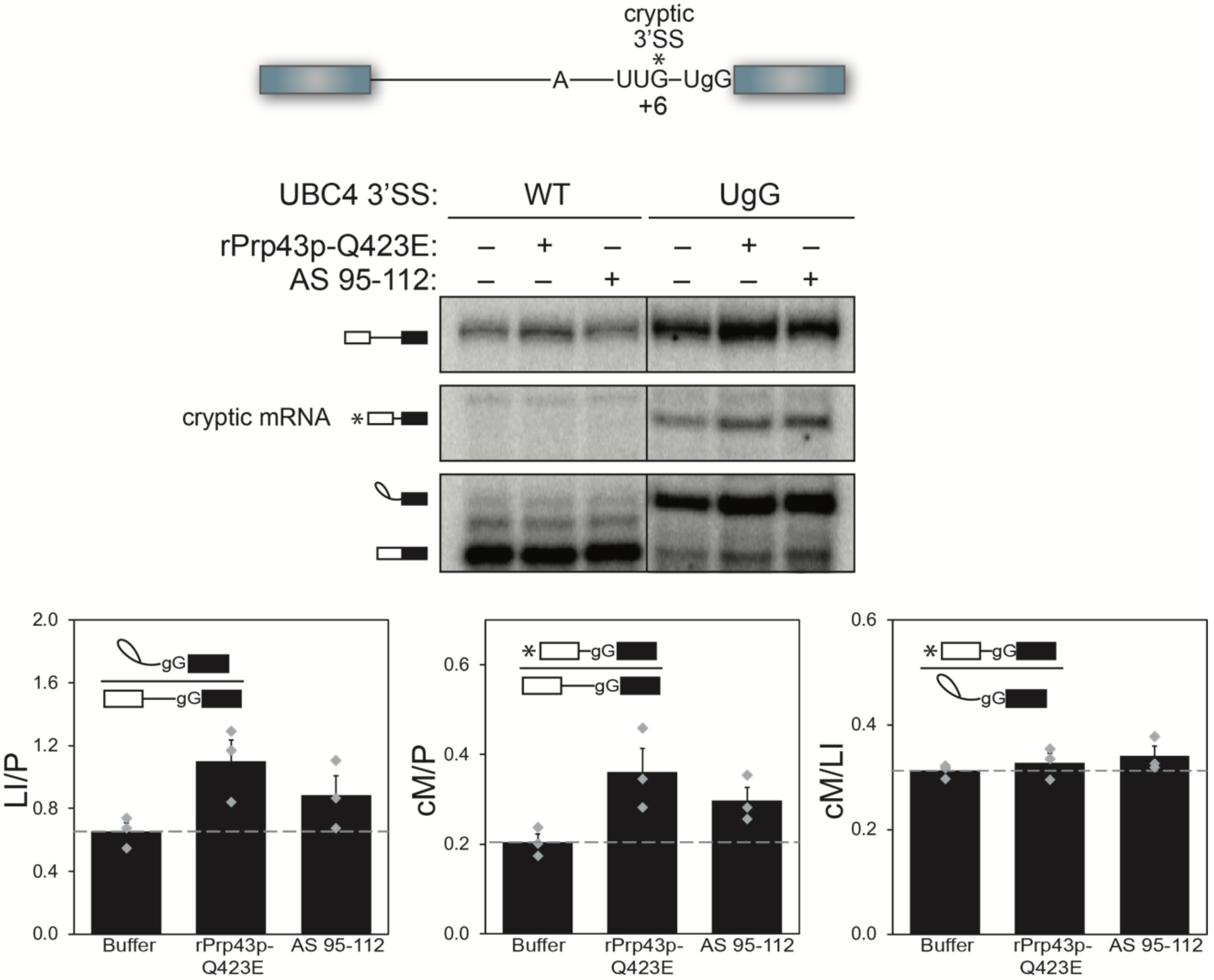

**<u>Figure S2</u>.** As for Prp43p (Mayas et al. 2010), the 3' end of U6 is required to minimize production of a cryptic mRNA. Unlabeled *UBC4* pre-mRNA with either a wild-type (WT) UAG 3’ splice site (SS) or mutated UgG 3' SS, as depicted in the schematic, was spliced in extract (ySCC1, (Chan et al. 2003)) that was first subjected to RNase H cleavage directed by DNA oligo AS 95-112 or that was supplemented with either buffer or rPrp43p-Q423E (Leeds et al. 2006). Reactions were then analyzed by extension of a radiolabeled 3' exon primer (oJPS239) and fractionation by denaturing PAGE. An asterisk indicates the cryptic 3’ splice site (UUG, located 6 nucleotides upstream of the mutated 3’ splice site) in the schematic or the corresponding, cryptic mRNA product in the gel image. Data are quantitated below with bars representing the mean $\pm$ SEM for three independent replicates, which are represented by grey diamonds. LI = lariat intermediate, P = pre-mRNA, cM = cryptic mRNA.

Toroney_FigS3

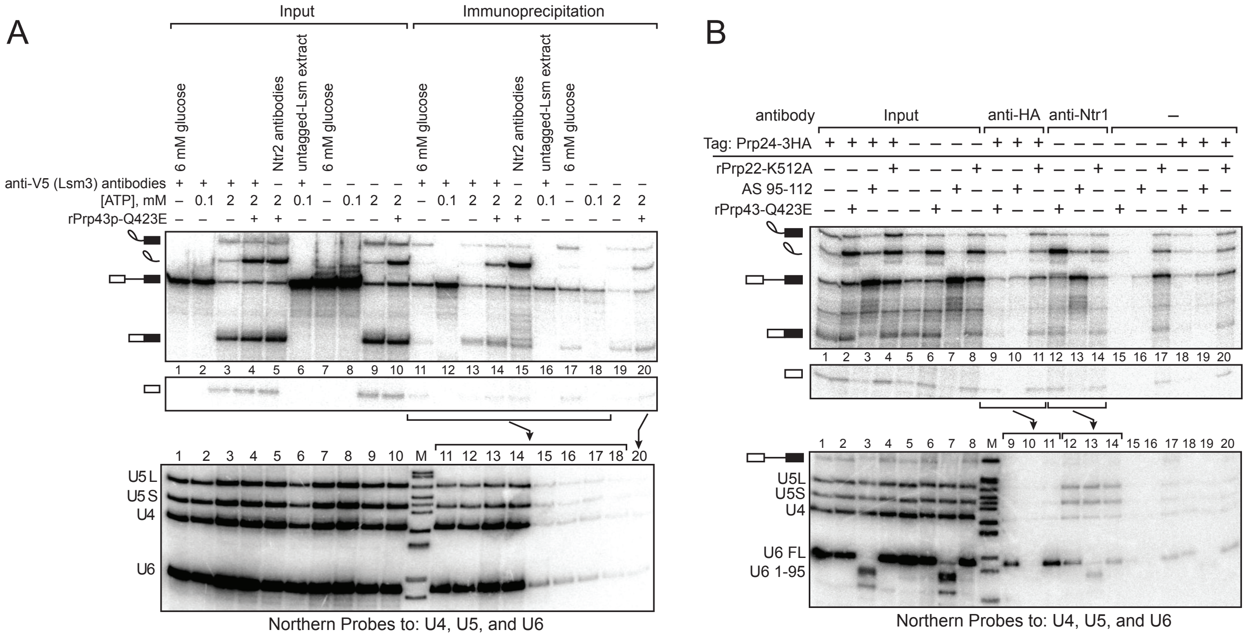

**<u>Figure S3</u>**. Lsm3p and Prp24p do not bind spliceosomes stalled at the disassembly stage. **A**. Lsm3p does not bind spliceosomes stalled at the disassembly stage. The top panel shows a denaturing PAGE analysis of *in vitro* splicing reactions assembled with radiolabeled *ACT1* pre-mRNA in V5-tagged- or untagged-Lsm3p yeast extracts (yJPS1448 or yJPS1405, respectively). Spliceosomes were immunoprecipitated with anti-V5 antibodies (lanes 11-14), anti-Ntr2p antibodies (lane 15) (Tanaka et al. 2007), or no antibodies (lanes 17-20). Twenty percent of each reaction was analyzed as input. As a positive control for pull-down of Lsm-bound spliceosomes, one reaction was carried out in the presence of low ATP (0.1 mM), which allows spliceosome assembly and thus binding of the Lsm complex but impedes activation and Lsm complex dissociation (Chan et al. 2003; Tarn et al. 1993); immunoprecipitation with anti-V5 antibodies in the presence of low ATP leads to enrichment of pre-mRNA as compared to the absence of ATP (cf. lanes 1-2 to 11-12) and as compared to the untagged control (cf. lanes 2 and 12 with lanes 6 and 16). To stall spliceosomes prior to release of excised intron, reactions were supplemented with rPrp43p-Q423E (Leeds et al. 2006), as indicated. In comparison with immunoprecipitation with anti-Ntr2p antibodies (lane 15), immunoprecipitation of tagged Lsm3p with anti-V5 antibodies did not result in significant enrichment of excised lariat intron relative to pre-mRNA or mRNA (cf. lanes 4 and 5 to lanes 14 and 15 as well as lane 20). Coimmunoprecipitation of snRNAs from the same reactions was monitored by northern blot (bottom panel) with radioactive probes directed to U5 (nts. 99-125), U4 (nts. 81-100), and U6 (nts. 28-54); as expected (Achsel et al. 1999; Gottschalk et al. 1999; Stevens and Abelson 1999), Lsm3p co-immunoprecipitated U4, U5, and U6, all components of the U4/U6.U5 tri-snRNP. **B**. Prp24p does not bind spliceosomes stalled at the disassembly stage. Denaturing PAGE analysis of *in vitro* splicing reactions using radiolabeled *ACT1* pre-mRNA in Prp24p-HA-tagged extract (yJPS867) or untagged extract (yJPS866) (Rader and Guthrie 2002). For the input lanes, 20% of each reaction was analyzed. Extracts were supplemented with buffer, mutated rPrp22p-K512A (Schwer and Gross 1998), or rPrp43p-Q423E (Leeds et al. 2006) or subjected to RNase H cleavage of U6 directed by DNA oligo AS 95-112. Reactions were immunoprecipitated with anti-HA 12CA5 antibodies, anti-Ntr1p antibodies (Tanaka et al. 2007), or no antibody. The reactions supplemented with mutated Prp22p, which stalls release of excised intron at a step prior to binding of NTR complex (Chen et al. 2013), served as a negative control, wherein the excised intron was not enriched by immunoprecipitation with anti-Ntr1p antibodies; by contrast, reactions supplemented with mutated Prp43p, which stalls intron released after NTR complex binding (Small et al. 2006), served as a positive control, wherein excised intron was enriched by immunoprecipitation with anti-Ntr1p antibodies (cf. input lanes 2 and 4 to lanes 12 and 14 and to lanes 15 and 17). For reactions stalled by mutated Prp43p, excised intron was effectively enriched by immunoprecipitation with anti-Ntr1p but not by antibodies against the HA tag on Prp24p (cf. input lanes 2 and 6 to lanes 9 and 12 and to lanes 15 and 18). This suggests that Prp24p does not associate with spliceosomes stalled just prior to disassembly. Coimmunoprecipitation of snRNAs was monitored by northern blot (bottom panel) with radioactive probes directed to U4, U5, and U6; the expected coimmunoprecipitation of U6 confirmed the functionality of the Prp24p tag and antibody (lanes 9 and 11)(Rader and Guthrie 2002; Shannon and Guthrie 1991). Note that Prp24p did not associate with free U6 when the last 10 nucleotides were trimmed by AS 95-112, as expected (cf. lane 10 to lanes 9 and 11) (Ryan et al. 2002).

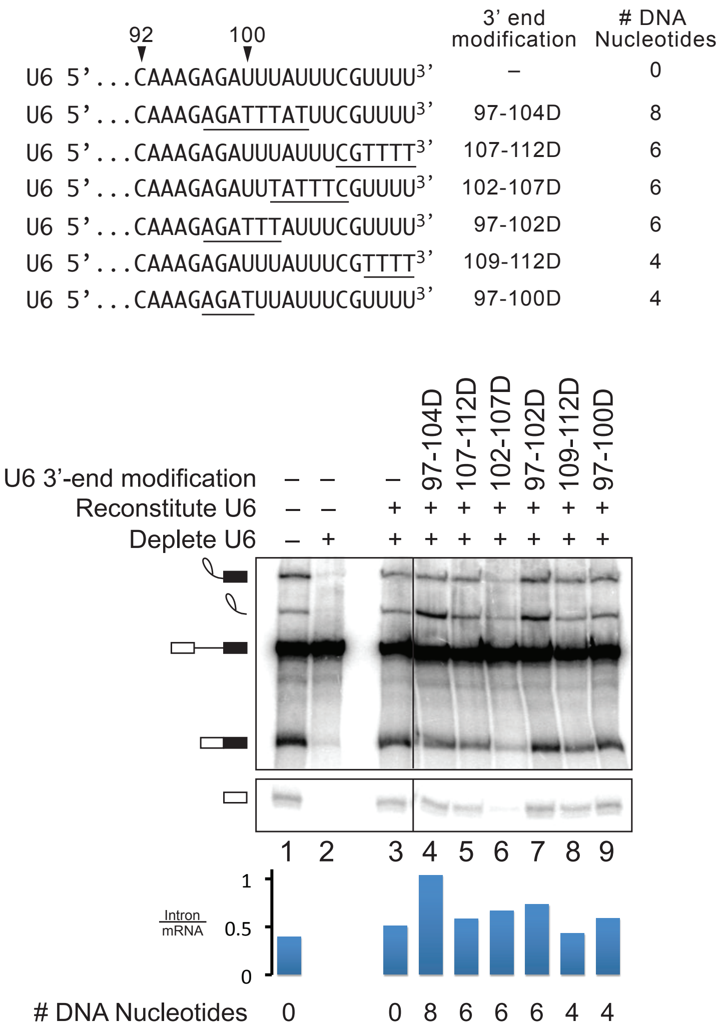


**<u>Figure S4</u>.** Excised intron release is inefficiently blocked by DNA substitutions of fewer than 8 nucleotides at the 3' end of U6. The sequence at the 3' end of U6 is depicted showing the position of deoxy substitutions (DNA bases are underlined) of either 8 nucleotides (97-104D), 6 nucleotides (107-112D, 102-107D, 97-102D) or 4 nucleotides (109-112D, 97-100D). Denaturing PAGE analysis is shown for *in vitro* splicing reactions using radiolabeled *ACT1* pre-mRNA in extracts (ySCC1) (Chan et al. 2003) reconstituted with the indicated U6 variants. Quantitation of intron turnover is shown below the gel.

Toroney_FigS5

A

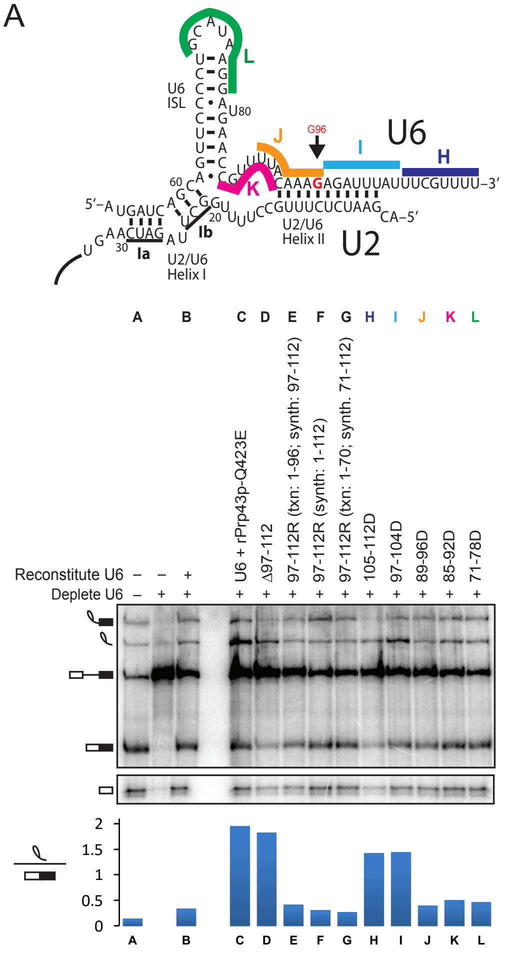


B

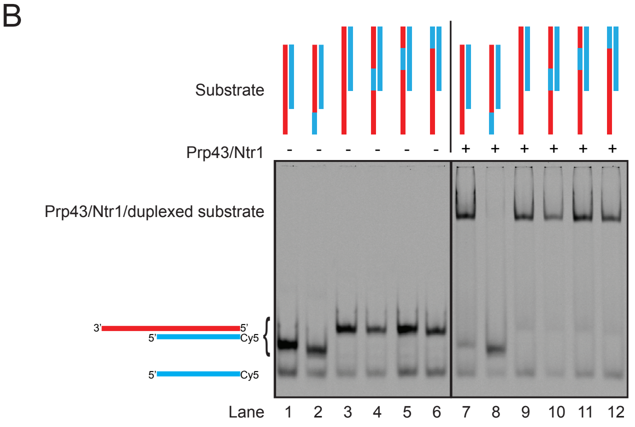

**<u>Figure S5</u>.** Excised intron release is not blocked by DNA substitutions in the U6 ISL or in the downstream bulge. **A.** DNA substitutions overlapping the loop of the U6 ISL or overlapping the bulge between the ISL and U2/U6 helix II do not impair intron turnover. Denaturing PAGE analysis of *in vitro* splicing reactions using radiolabeled *ACT1* pre-mRNA in extracts (ySCC1) (Chan et al. 2003) reconstituted with U6 variants illustrated in the U2/U6 schematic, wherein bars indicate positions of DNA substitutions within U6 (reactions H-L). Reactions A-G correspond to controls and include reactions reconstituted with full-length U6 and supplemented with mutated rPrp43-Q423E (**C**), reconstituted with truncated U6 (Δ97-112; **D**), or reconstituted with full-length U6 constructed from various segments of transcribed ("txn") or synthetic ("synth") RNA (**E-G**). Intron turnover was quantitated as the ratio of excised intron to mRNA and is shown below the gel. **B.** rPrp43p/Ntr1p binds to all unwinding substrates tested in Figs. 5G and 5H except the substrate with a DNA substitution in the single stranded overhang (cf. lanes 2 and 8), which was analyzed for unwinding in Fig. 5G (lanes 6-10). The migration of substrates was analyzed after incubation without (-) or with (+) rPrp43p/Ntrp1(1-120). The substrates are cartooned above each lane and coded as in Fig. 5G and 5H, with red indicating RNA and blue indicating DNA. Note that low levels of single-stranded, labeled oligo contaminated the preparations of duplexed substrate. The vertical dividing line indicates that lanes containing substrates not described in the manuscript were omitted for clarity.

Toroney_FigS6

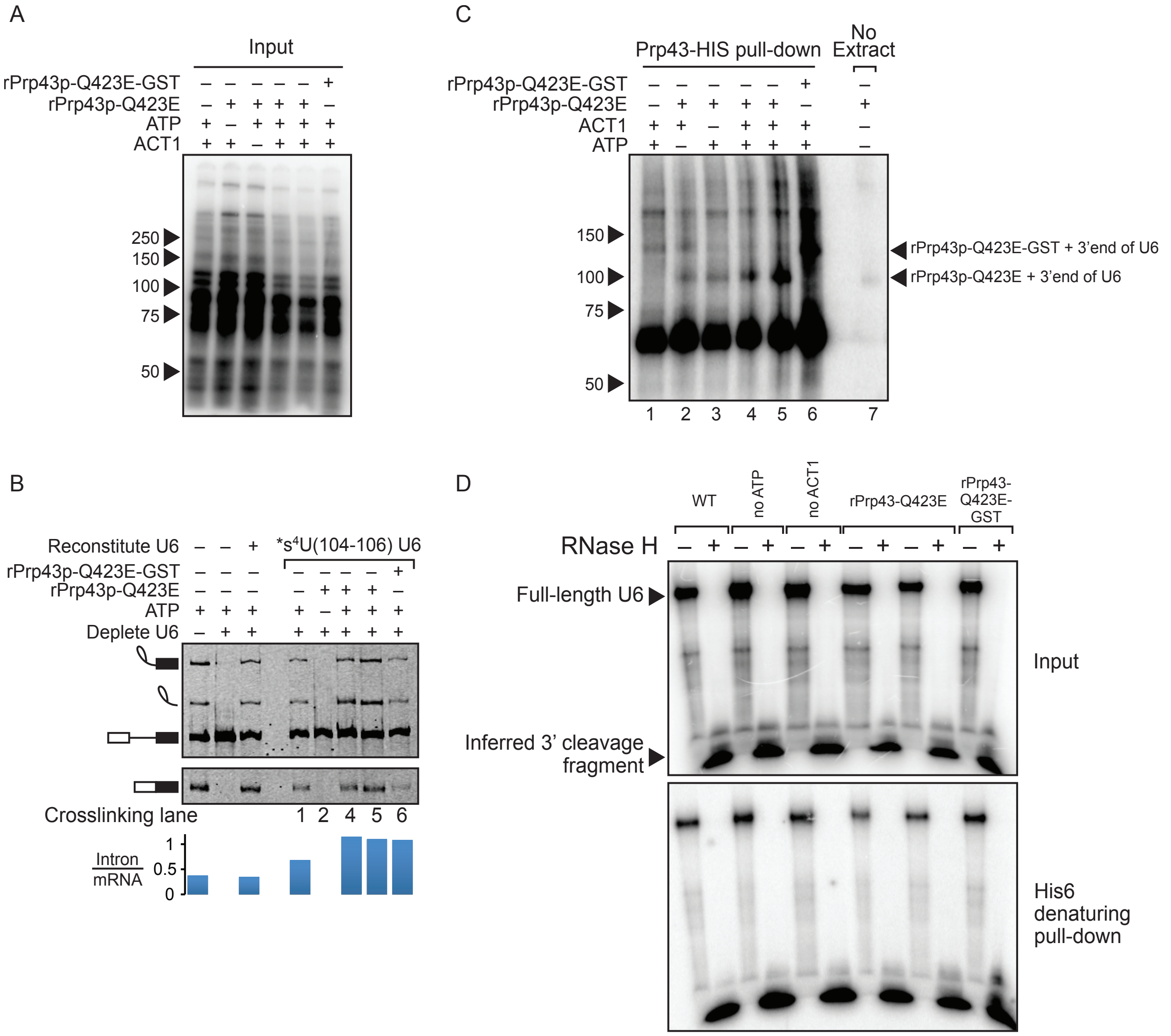

**<u>Figure S6</u>.** A UV crosslink at the 3' end of U6 snRNA can be immunoprecipitated with Prp43p under denaturing conditions. **A**. SDS-PAGE analysis of crosslinked reactions used as inputs for immunoprecipitation experiments described in Figures 6B and S6C, prior to RNase H digestion. **B**. Denaturing PAGE analysis of crosslinked reactions of Cy5-*ACT1* splicing described in Figures 6B and S6C. Intron turnover is quantified below the gel. **C**. SDS-PAGE analysis of crosslinked splicing reactions performed in extracts reconstituted with *$s^4$U(104-106) U6 and supplemented where indicated with either rPrp43p-Q423E or GST-tagged rPrp43p-Q423E (Leeds et al. 2006). A parallel reaction was performed in the absence of extract but with rPrp43p-Q423E as indicated (lane 7). The reactions were subjected to denaturing conditions under which the His6-tag on the recombinant proteins was used to pull-down cross-linked species using Ni-NTA beads. Immunoprecipitated reactions were then subjected to RNase H digestion of U6 directed by the DNA oligo AS 48-93, as well as by the DNA oligo AS 28-54 to ensure efficient cleavage of U6 at the 3' end via AS 48-93. The migration of protein markers is indicated to the left of the gel, and the expected position of RNase H-digested U6 crosslinked to rPrp43-Q423E or GST-tagged rPrp43p-Q423E is indicated to the right. Note that the identity of the prominent band at ~65 kDa is unknown, but the band is enriched after Ni-NTA pull-down down independent of added recombinant, His6-tagged Prp43p. **D**. Denaturing PAGE analysis of radiolabeled *$s^4$U(104-106) U6 extracted from reactions analyzed in panels A, B, and C before and after RNase H digestion. The panel shows that cleavage of U6 was efficient and that the inferred 3' cleavage fragment pulled-down with Prp43p via the His6 tag.

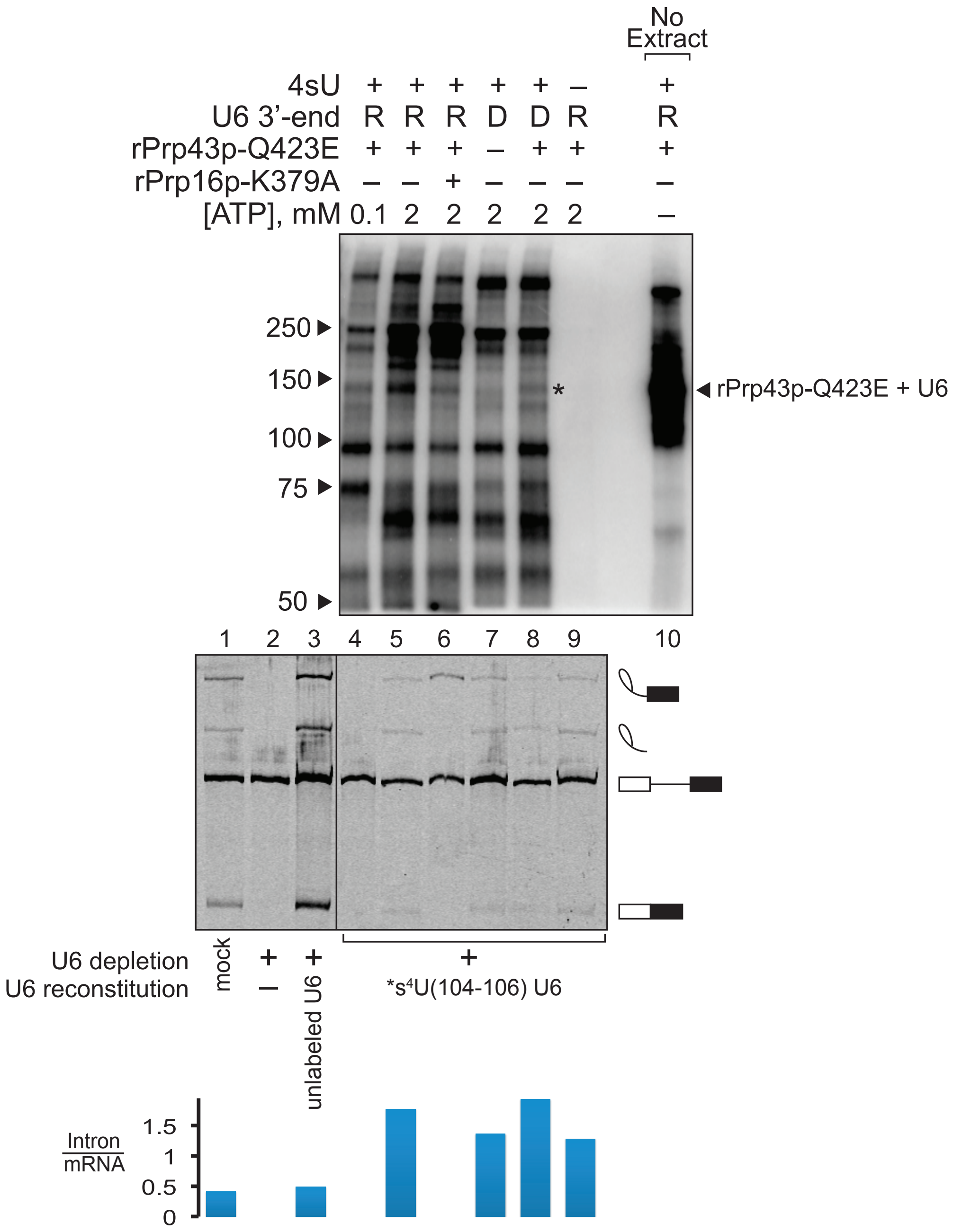

No Extract
4sU + + + + + – +
U6 3'-end R R R D D R R
rPrp43p-Q423E + + + – + + +
rPrp16p-K379A – – + – – – –
[ATP], mM 0.1 2 2 2 2 2 –
250
150
100
75
50
*
rPrp43p-Q423E + U6
1 2 3 4 5 6 7 8 9 10
U6 depletion
U6 reconstitution
mock
+ –
+ unlabeled U6
+
*s4U(104-106) U6
Intron/mRNA
1.5
1
0.5
0

**<u>Figure S7</u>.** A UV crosslink at the 3' end of U6 with Prp43p is dependent on $s^4U$, as well as on ATP, Prp16p, and RNA at the 3' end. SDS-PAGE analysis of crosslinked splicing reactions performed in extracts reconstituted with U6 containing a single radiolabel between G96 and A97 but lacking $s^4U$ substitutions (lane 9), compared with reactions reconstituted with *$s^4U$(104-106) U6 (lanes 4-6) and *$s^4U$(104-106) U6 containing deoxy substitutions ("D") at nucleotides 97-103 and 107-112 (lanes 7 and 8). Reactions were supplemented where indicated with rPrp16p-K379A (Schneider et al. 2002) or with rPrp43-Q423E (Leeds et al. 2006) in the presence of high (2 mM) or low (0.1 mM) ATP. Activated spliceosomes in the reactions were immunoprecipitated under native conditions via HA-tagged Prp19p. The asterisk indicates migration of rPrp43-Q423E crosslinked to U6 in the absence of extract (lane 10). Splicing of Cy5-*ACT1* was monitored by denaturing PAGE (bottom panel), and intron turnover was quantified below; as expected, low ATP precluded splicing (Tarn et al. 1993); mutated Prp16p blocked exon ligation (Schneider et al. 2002); and mutated Prp43p blocked spliceosome disassembly (Tsai et al. 2007; Small et al. 2006; Martin et al. 2002), as reflected by the increase in the excised intron:mRNA ratio.
